# Histone modifications regulate pioneer transcription factor binding and cooperativity

**DOI:** 10.1101/2023.03.14.532583

**Authors:** Kalyan Sinha, Silvija Bilokapic, Yongming Du, Deepshikha Malik, Mario Halic

## Abstract

Pioneer transcription factors have the ability to access DNA in compacted chromatin. Multiple transcription factors can bind together to a regulatory element in a cooperative way and cooperation between pioneer transcription factors Oct4 and Sox2 is important for pluripotency and reprogramming. However, the molecular mechanisms by which pioneer transcription factors function and cooperate remain unclear. Here we present cryo-EM structures of human Oct4 bound to a nucleosome containing human Lin28B and nMatn1 DNA sequences, which bear multiple binding sites for Oct4. Our structural and biochemistry data reveal that Oct4 binding induces changes to the nucleosome structure, repositions the nucleosomal DNA and facilitates cooperative binding of additional Oct4 and of Sox2 to their internal binding sites. The flexible activation domain of Oct4 contacts the histone H4 N-terminal tail, altering its conformation and thus promoting chromatin decompaction. Moreover, the DNA binding domain of Oct4 engages with histone H3 N-terminal tail, and posttranslational modifications at H3K27 modulate DNA positioning and affect transcription factor cooperativity. Thus, our data show that the epigenetic landscape can regulate Oct4 activity to ensure proper cell reprogramming.

## Introduction

DNA-binding transcription factors (TFs) target distinct DNA sequences at gene regulatory regions, thus ensuring specificity in transcription machinery assembly ^1, 2^. DNA packaging into nucleosomes can hinder TF binding to target sequences ^3^, but a small set of so-called pioneer TFs can access DNA even within compacted chromatin ^4–8^. Once bound to their target sites, pioneer TFs can facilitate the recruitment of other TFs by creating accessible chromatin, a property that underlies their function as master regulators in embryo development, cell differentiation and reprogramming. In fact, over-expression of four pioneer TFs — Oct4, Sox2, Klf4 and c-Myc — promotes the reprogramming of cells to pluripotency ^9^, with Oct4 expression being necessary and sufficient to reprogram cells ^10, 11^. *In vitro* Sox2, Klf4 and c-Myc bind to nucleosome more efficiently in presence of Oct4 ^6^ and cooperativity between Oct4 and Sox2 is critical for early development and reprogramming ^12–18^, but the molecular mechanisms involved remain unclear.

Oct4 has two DNA-binding domains, called Oct4_POU_S_ and Oct4_POU_HD_. Previous X-ray structures showed the two domains wrapping around naked DNA, but such binding mode would be incompatible with the nucleosome architecture ^19, 20^. In recent cryo-EM work ^21^, Oct4_POU_S_ and the DNA-binding domain of Sox2 were seen unwrapping a nucleosome containing binding sites for those TFs inserted into the DNA positioning sequence 601 ^22^. The inserts were placed to promote optimal binding and stability of the complex, but the 601 sequence is known to suppress the nucleosome dynamics that is typical of biologically relevant sequences ^22^. In recent efforts to capture Oct4 bound to a nucleosome with an endogenous DNA sequence, the density for Oct4 could not be observed ^23, 24^. Hence, a structure of Oct4 in complex with a nucleosome containing a physiologically relevant DNA sequence remained elusive, limiting our mechanistic understanding of pioneer TF function. For instance, although Oct4 and other TFs bind nucleosomes, it remains unclear whether they interact with histones and whether epigenetic marks would affect that interaction. Previous crosslinking/mass spectrometry analyses of reconstituted Oct4–nucleosome complex with endogenous DNA have shown that Oct4 binds near histone H3 N-terminal tail ^24^. Direct interactions between TF and histones would require proper positioning of the DNA binding site on the nucleosome, which could be hindered by the use of artificial DNA sequences such as 601. Moreover, chromatin occupancy by Oct4 correlates with histone marks found in enhancers, such as H3K27ac and H3K4me1, but silent marks such as H3K27me3 are also found at Oct4 binding sites ^25–30^; it remains to be determined whether and how those modifications regulate Oct4 binding.

To address these gaps, we present cryo-EM structures of human Oct4 bound to nucleosomes containing DNA sequences from human Lin28B or nMatn1 loci, along with biochemistry assays. The Lin28B sequence has 3 binding sites for Oct4, as well as binding sites for Sox2, Klf4 and c-Myc ^6^, whereas nMatn1 has multiple Oct4 binding sites. Both sequences are thus an ideal platform to study cooperative assembly of multiple pioneer TFs. We find that on both nucleosomes, Lin28B and nMatn1, Oct4 engages a binding site that is transiently exposed through DNA sliding. Binding of that first Oct4 molecule stabilizes the positioning of DNA on the nucleosome, to allow binding of additional Oct4 and of Sox2 to their internal binding sites, explaining the basis of pioneer transcription factor cooperativity. Remarkably, the structures reveal direct contacts between Oct4 and histones H2A, H3 and H4, providing direct evidence that TFs do not interact solely with their target DNA. The intrinsically disordered activation domain of Oct4 binds and remodels histone H4 tail, thus regulating chromatin decompaction. Furthermore, we show that H3K27 modifications can regulate Oct4 interactions with the nucleosome and affect DNA positioning, which in turn determines cooperative TF binding. Together, our observations reveal that Oct4 binds histones and that histone modifications regulate cooperative binding of pioneer TFs.

## Results

### Oct4 binding positions Lin28B nucleosome

To investigate the mechanism for cooperativity between pioneer TFs Oct4 and Sox2, we assembled a complex containing full-length human Oct4 and Sox2 and a nucleosome with DNA from the human Lin28B locus ^31–33^. This DNA fragment contains three binding sites for Oct4 (binding sites 1, 2 and 3, or OBS1–3) and one for Sox2 ^6^. By native gel electrophoresis assays, we observed association of Oct4 and Sox2 with nucleosomes assembled with Lin28B DNA fragments that were 162- or 182-bp long (**Extended Data Fig. 1a**). However, by cryo-EM analyses, we could only visualize the proteins bound to the 182-bp nucleosome, indicating that the complex on the nucleosome with shorter DNA is less stable ^24^. Hence, for the remainder of this work, we used exclusively the 182-bp nucleosome.

The initial cryo-EM reconstructions showed a density bound to the linker DNA (**Extended Data Fig. 1b-h, Table 1**), but the resolution was limited because of flexibility of the complex. To improve the resolution, we used focused classification followed by local search refinements and obtained maps with resolutions of 2.8 Å in the nucleosome portion (**Extended Data Fig. 1g-i**) and of 3.9 Å for a 30-kDa region of Oct4 bound to linker DNA (**Extended Data Fig. 2a-g**). We did not observe clear density for Sox2, suggesting that it might have dissociated during sample preparation. The two maps had sufficient overlapping densities to allow assembly of a composite map and model (**Fig. 1a, Extended Data Fig. 2h, Table 1**). In the structure, Oct4 is bound to the linker DNA near the nucleosome entry/exit site (**Fig. 1a, Extended Data Fig. 2h**), in agreement with previous crosslinking mapping of reconstituted complexes ^24^. Notably, the DNA bases are well resolved along the nucleosome wrapped region, indicating minimal movement of the Lin28B sequence in the complex with Oct4 (**Extended Data Fig. 1i**). The high resolution of the nucleosomal DNA allowed us to precisely position the sequence, with the Oct4 binding site 1 (OBS1) placed at the exact location of the Oct4 density (**Fig. 1a, Extended Data Fig. 1i, 2i**).

**Fig. 1.**
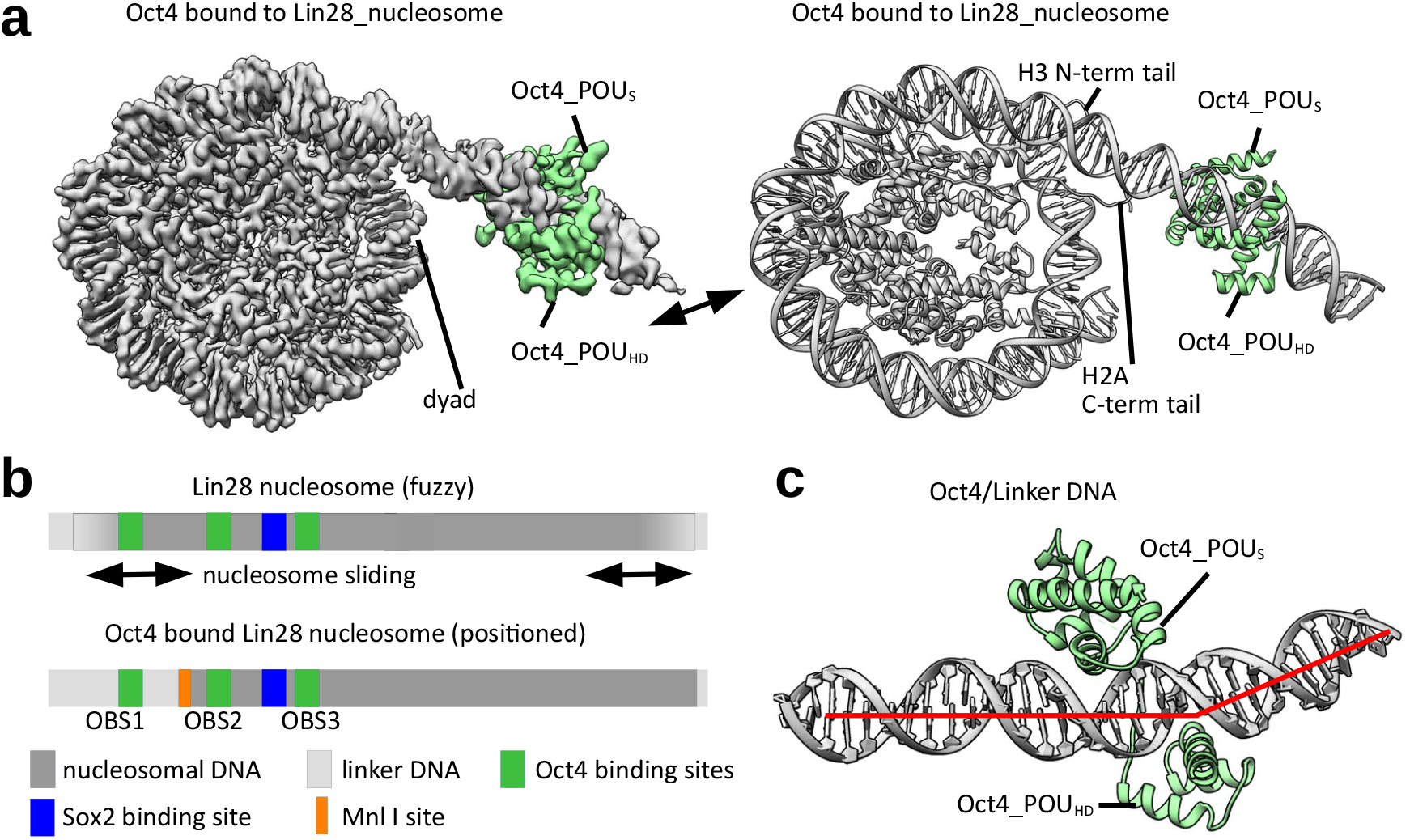
Oct4 binds nucleosome at exposed DNA site. **a)** A composite cryo-EM map (left) and structural model (right) of human Oct4 (green) bound to a nucleosome (grey) assembled with a 182-bp DNA fragment from the Lin28B locus. **b)** Schematic representation of DNA positioning on the Lin28B nucleosome. Binding sites for Oct4 (OBS1, 2, 3) and Sox2, and Mnl1 restriction site are shown. Top, the nucleosome is “fuzzy” as the DNA adopts multiple positions represented by bidirectional arrows. Bottom, Oct4 binding stabilizes DNA at a defined position on the nucleosome. **c)** Close-up view of Oct4 (green) bound to the nucleosomal DNA (grey; red line shows the path of the DNA helix axis), showing the kink in the linker DNA introduced by Oct4_POU_HD_.

To determine if Oct4 stabilizes DNA positioning on the Lin28B nucleosome, we determined a cryo-EM structure of that same nucleosome in absence of any TFs (**Table 1**). This structure had an overall resolution of 3.1 Å, similar to the Oct4-bound nucleosome, but the DNA bases were not well resolved and hence the DNA position could not be determined (**Extended Data Fig. 3a-f**). We also observed that in some particle classes, the linker DNA protrudes from both sides of the histone octamer (**Extended Data Fig. 3g**), whereas in the Oct4-bound structure, it protrudes from only one side. Together, these observations indicate that the Lin28B DNA can adopt several positions on the free nucleosome (**Extended Data Fig. 3g**), in contrast to its well-defined positioning in the Oct4-bound nucleosome (**Fig. 1a**). These findings are consistent with *in vivo* data showing that the nucleosome at the Lin28B locus is “fuzzy” and occupies ~200 bp ^34^. Thus, the naturally occurring Lin28B sequence is able to move along the histone octamer, transiently exposing OBS1. Once Oct4 binds OBS1, it traps the DNA in that position and stabilizes the otherwise flexible linker DNA into a more defined conformation (**Fig. 1b, Extended Data Fig. 3h**). Notably, we observed that hexasomes are 3-fold more abundant in Oct4-bound sample when compared to the free Lin28B nucleosomes (**Extended Data Fig. 1g, 3f**).

In the structure, both DNA-binding domains of Oct4 engage with the Lin28B nucleosome: Oct4_POU_S_ is bound to the linker DNA, close to the nucleosome dyad, whereas Oct4_POU_HD_ is located distally from the nucleosome (**Fig. 1a, Extended Data Fig. 2h**). These observations are consistent with the requirement for both Oct4 DNA-binding domains to efficiently bind chromatin *in vivo* ^21^ (**Fig. 1a, c**). The Oct4 interactions with DNA in our structure are overall similar to those observed in the crystal structure of Oct4 bound to naked DNA ^19^, but we see that Oct4_POU_HD_ introduces a kink in the linker DNA, due to arginine residues widening the DNA major groove (**Fig. 1c, Extended Data Fig. 2g, 3i, j**); such DNA distortion could disrupt local chromatin organization and affect binding of other proteins. Overall, the Oct4-DNA interactions in our structure differ considerably from those seen in the cryo-EM structure of Oct4 bound to a 601-based nucleosome, in which only Oct4_POU_S_ was observed to interact with the nucleosome ^21^.

### Oct4 N-terminal region changes the H4 tail conformation

Having shown that Oct4 binding stabilizes the positioning of nucleosomal DNA, we examined whether it induced other changes to the nucleosome structure. We observed in our cryo-EM data that the H4 N-terminal tail on the Oct4-proximal side of the nucleosome adopts multiple conformations, while the H4 tail on the opposite side is predominately found in the canonical conformation, following the DNA path at SHL 2 ^35, 36^ (**Extended Data Fig. 4a**). Further image classification revealed two major conformations for the H4 tail on the Oct4-proximal side of nucleosome: the first one resembles the canonical conformation, whereas in the second one, the H4 tail is rotated 90° towards SHL 1 and an additional density can be seen interacting with the H4 tail and α2 helix (**Fig. 2a, Extended Data Fig. 4a-c**). This density was not observed on the Lin28B nucleosome or on the Oct4-distal side of the Oct4-bound nucleosome, and we hypothesized that it could originate from the Oct4 activation domain, which consists of flexible N- and C-terminal regions. The density alters the conformation of Asp24 at the beginning of the H4 tail, which in turn changes the conformation of the whole H4 tail (**Fig. 2a**), moving residues that are essential for chromatin compaction by more than 30 Å and potentially disrupting interactions between nucleosomes.

**Fig. 2.**
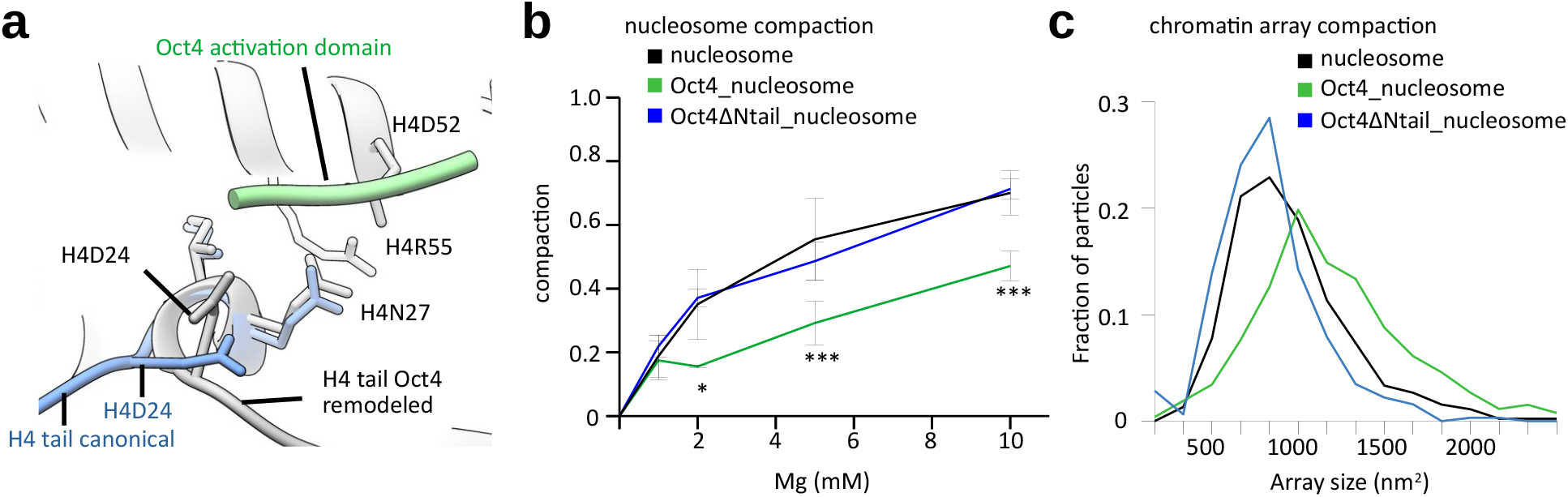
Oct4 activation domain remodels the H4 N-terminal tail to open chromatin. **a)** Overlay of the two distinct conformations of the H4 tail on the Oct4-proximal side of the nucleosome: canonical conformation (blue) and Oct4-remodeled conformation (grey). Interaction of the disordered region of Oct4 (green) with the H4 tail rearranges Asp24, leading to repositioning of the whole tail. **b)** Mono-nucleosome compaction by Mg^2+^, assessed by native gel electrophoresis; a representative gel is shown in Extended Data 5a. Compaction was quantified by the reduction in the intensity of the nucleosome band, due to nucleosome precipitation. Data shown are mean and s.e.m. of 4 independent measurements. * p<0.05, ** p<0.01, *** p<0.001, Student’s t-test comparing Oct4 bound nucleosome and nucleosome. **c)** Quantification of negative-stain EM data showing Mg^2+^-induced compaction of nucleosome arrays assembled on a 1022-bp-long DNA fragment from the Lin28B locus. Graph shows distribution of the area occupied (size) by nucleosome arrays, measured from micrographs with the different samples; 300-450 arrays were analyzed per sample. Representative micrographs are shown in Extended Data Fig. 5c.

These observations suggest that Oct4 binding might affect the interactions between two nucleosomes. Further analyses of our cryo-EM data for interacting nucleosomes ^37^ showed that, in the absence of Oct4, 9% of nucleosomes interact on the cryo EM grid, predominantly near the linker DNA (**Extended Data Fig. 4d**). This interaction mode is abolished and the overall frequency of nucleosome interactions was lower in the Oct4-bound subset, with 4.2% of nucleosome engaging in two types of arrangements (**Extended Data Fig. 4d**): on the Oct4 distal side (side B), two nucleosomes stack against each other mostly on the histone octamer side, as previously observed ^37^ (2.7% of particles). In contrast, on the Oct4 proximal side, the second nucleosome is pushed away from Oct4 to the back of the first nucleosome (1.5% of particles), leading to a more open chromatin architecture. These data suggest that the presence of bound Oct4 and the alternative H4 tail conformation on the Oct4-proximal side reduce interaction with another nucleosome. To test this hypothesis, we induced nucleosome compaction with Mg^2+^ ^38, 39^ and found that the presence of Oct4 substantially reduced association between mononucleosomes as assessed by native gel electrophoresis (**Fig. 2b, Extended Data Fig. 5a**). We also assembled chromatin arrays using a longer 1040 bp DNA fragment from the Lin28B locus and examined their compaction with Mg^2+^ by negative-stain EM imaging: we found that the nucleosome arrays are more open in the presence of Oct4 (**Fig. 2c, Extended Data Fig. 5b-c**).

Finally, we tested the roles of the N- and C-terminal flexible regions of Oct4 on chromatin decompaction by deleting them individually. Neither deletion reduced the interaction of Oct4 with the Lin28B nucleosome (**Extended Data Fig. 5d**). However, Oct4 lacking the N-terminal region lost the ability to reduce nucleosome compaction and inter-nucleosome interactions, whereas deletion of the C-terminal tail did not affect those properties (**Fig. 2b, c, Extended Data Fig. 5e**). Together, our structural and biochemical data suggest that the N-terminal region of Oct4 remodels the H4 tail and contributes to chromatin decompaction.

### Oct4 cooperativity is modulated by histone marks

In our structure of Oct4-bound nucleosome, both the Oct4-binding sites OBS2 and OBS3 have partial internal motifs exposed, which would allow binding of Oct4_POU_HD_ to OBS2 and Oct4_POU_S_ to OBS3 (**Fig. 3a**). Binding to partial DNA motifs has been previously proposed ^6^ and our data suggests that DNA positioning induced by binding of Oct4 to OBS1 facilitates binding of additional Oct4 molecules to their internal sites.

**Fig. 3.**
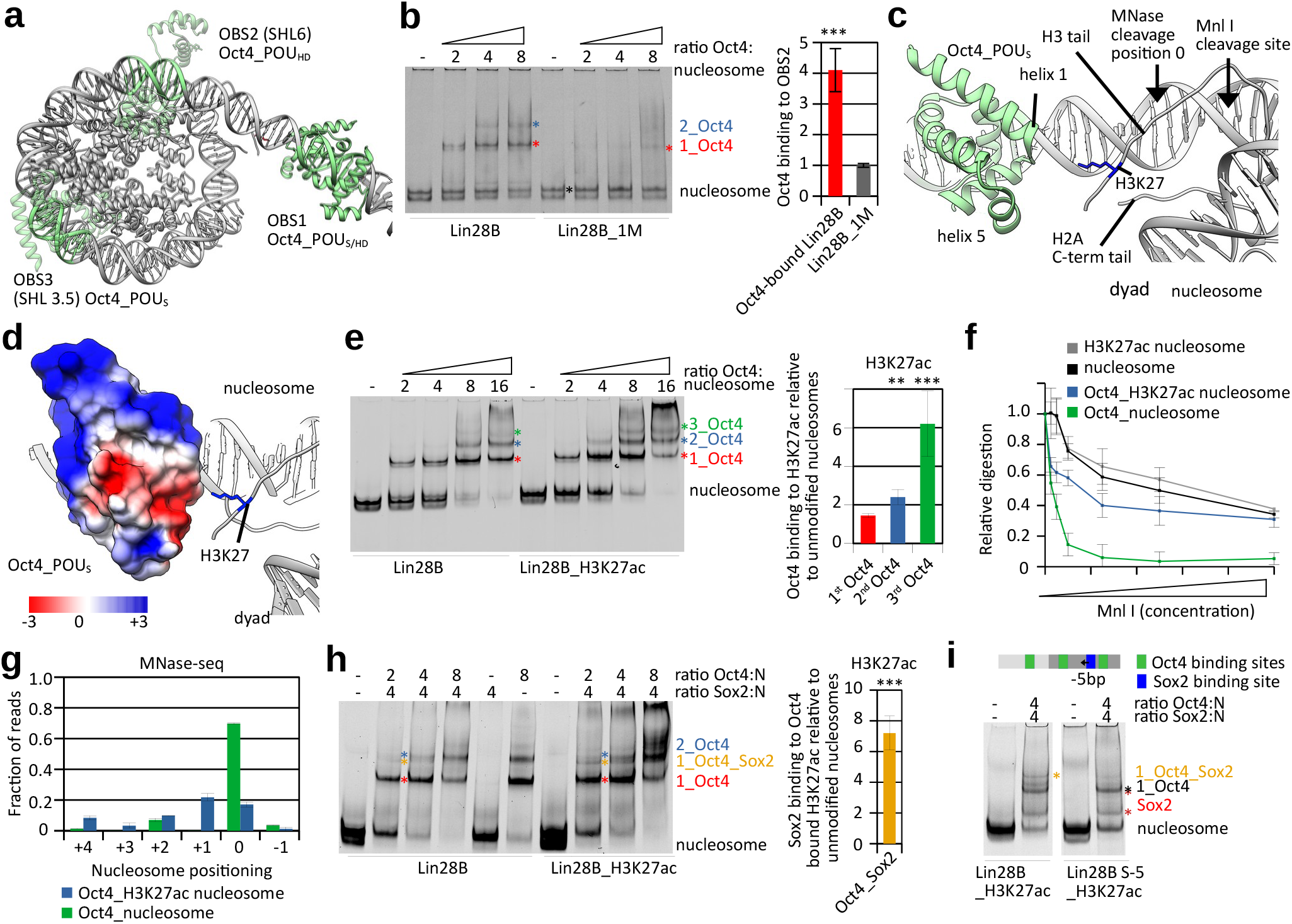
Histone modifications modulate Oct4 and Sox2 cooperativity. **a)** Model of Oct4 bound to the Lin28B nucleosome, showing Oct4-binding sites on DNA in green. Oct4 bound to OBS1 is in solid green; Oct4 structure (transparent green) was superimposed on OBS2 and on OBS3. Oct4_POU_HD_ can bind to OBS2 and Oct4_POU_S_ can bind to OBS3. **b)** Left, representative native gel electrophoresis showing Oct4 binding to Lin28B or Lin28B_1M (with mutated OBS1) nucleosomes. The composition of the Oct-bound bands was validated by western blot analyses (Extended Data Fig. 5g). Colored asterisks mark the number of Oct4 molecules bound to the nucleosome: black, nucleosome alone; red, 1 Oct4 bound; blue, 2 Oct4 bound. For Lin28B samples, compare 2_Oct4 band (blue asterisk) to 1_Oct4 band (1 Oct4 bound to OBS1, red asterisk). For Lin28B_1M samples, compare 1_Oct4 band (nucleosome with 1 Oct4 bound to either OBS2 or 3, red asterisk) to input nucleosome (black asterisk). Right, quantification of the native gel electrophoresis data, using bands marked with asterisks and showing the described comparisons as ratios. Data are mean and s.e.m. of 4 independent experiments; *** p<0.001, Student’s t-test. **c)** Close-up views of the nucleosome entry/exit site showing interaction of the Oct4_POU_S_ domain with the H3 N-terminal and H2A C-terminal tails near the dyad. Ribbon representation shows Oct4_POU_S_ helix 1 and helix 5 interacting with histone H3 N-terminal tail; H3K27 is shown in blue. **d)** Surface model showing electrostatic potential of Oct4_POU_S_ domain. The positively charged H3 tail is near a negatively charged surface of Oct4. **e)** Left, representative native gel electrophoresis showing Oct4 binding to the Lin28B or Lin28B_H3K27ac nucleosomes. The composition of the Oct4-bound bands was validated by western blot analyses (Extended Data Fig. 5g). Colored asterisks indicate the number of Oct4 molecules bound to the nucleosome: red, 1 Oct4; blue, 2 Oct4; and green, 3 Oct4. Right, quantification of the native gel electrophoresis data, using bands marked with asterisks. Data are mean and s.e.m. of 4 independent experiments; ** p<0.01, *** p<0.001, Student’s t-test. **f)** Quantification of Mnl I digestion of free and Oct4-bound nucleosomes, unmodified or with H3K27ac. The y-axis shows intensity of the intact Oct4_nucleosome complex or nucleosome band after enzyme digestion, normalized to the input (without enzyme). Data are mean and s.e.m. of 4 independent experiments. Representative gels are shown in Extended Data Fig. 6d. **g)** Quantification of sequencing of MNase I-digested Oct4-bound nucleosomes, unmodified or with H3K27ac. The y-axis shows fraction of nucleosome size reads starting at defined position, the x-axis shows position of the first base pair relative to the most abundant position (0 as observed in the structure). Data are mean and s.e.m. of 2 independent experiments. A more detailed representation is shown in Extended Data Fig. 6f. **h)** Left, representative native gel electrophoresis showing Oct4 and Sox2 binding to Lin28B and Lin28B_H3K27ac nucleosomes. Sox2 concentration is constant, whereas Oct4 concentration is increased as indicated. Colored asterisks indicate nucleosomes with different molecules bound: red, 1 Oct4; orange, 1 Oct4 and 1 Sox2; blue, 2 Oct4. Compare 1_Oct4_Sox2 band (orange asterisk) to 1_Oct4 band (red asterisk) in Lin28B and Lin28B_H3K27ac nucleosomes. Right, quantification of the native gel electrophoresis, using bands marked with asterisks. Data shown as mean and s.e.m. of 6 independent experiments; *** p<0.001, Student’s t-test. **i)** Left, representative native gel electrophoresis showing Oct4 and Sox2 binding to Lin28B_H3K27ac nucleosomes and Lin28B_H3K27ac with Sox2 binding site moved for 5bp. Repositioning of the binding site does not affect Sox2 binding to nucleosome (grey asterisk), but strongly reduces cooperative binding of Oct4 and Sox2 (orange asterisk). Lower red asterisk shows Sox2 degradation product (Sox2 DBD) bound to nucleosome.

To test this hypothesis, we mutated each of the Oct4-binding sites in Lin28B sequence and examined binding of Oct4 to nucleosomes by native gel electrophoresis. This setup allows us to distinguish nucleosomes with one or more Oct4 bound (**Extended Data Fig. 5f**). With the wild type Lin28B nucleosome, we can detect strong binding of one Oct4 but also binding of a second and weaker binding of the third Oct4 **(Extended Data Fig. 5f, g**). Mutation of either OBS2 or OBS3 did not affect formation of a complex with one Oct4 bound, indicating that the first Oct4 binds to OBS1 as in the wild-type Lin28B nucleosome, whereas a second Oct4 binds either OBS2 or OBS3 (**Extended Data Fig. 5h, i**). In contrast, when we mutated OBS1 to generate Lin28B_1M nucleosome, Oct4 binding to OBS2/3 was considerably reduced compared to the wild-type Lin28B nucleosome or mutated on OBS2/3 (**Fig. 3b**). Thus, Oct4 binding to OBS2/3 is stimulated when a first Oct4 is bound to OBS1, which is in agreement with our structural data showing that binding of Oct4 to OBS1 stabilizes nucleosomal DNA and exposes partial motifs in internal OBS2/3 sites.

We turned our attention back to Oct4 DNA-binding domains bound to OBS1, and observed interactions between Oct4_POU_S_ and histones H3 and H2A: the tip of helix 1 (residues 159–163) contacts histone H2A C-terminal tail and histone H3 N-terminal tail; the latter is also contacted by small helix 5 (residues 213–222) (**Fig. 3c and Extended Data Fig. 5j**). The dipole moment of helix 1 and negatively charged helix 5 together form an acidic patch on Oct4 that faces the nucleosomal dyad and interacts with positively charged histone tails there, mediating additional interaction between Oct4 DNA binding domain and nucleosome (**Fig. 3d**).

In our structure of Oct4-bound nucleosome, histone H3K27 is in close proximity to the acidic patch of the Oct4, specifically to the tip of helix 1, suggesting a potential electrostatic interaction between positively charged H3K27 and negative dipole moment of helix 1 (**Fig. 3c, d**). This observation prompted us to examine whether H3K27 modifications would affect Oct4 binding to the nucleosome. H3K27 acetylation (H3K27ac) is an active mark associated with enhancers that was found to colocalize with Oct4 on chromatin ^26–28^. Acetylation of H3K27 would neutralize the positive charge of the Lys residue and would be expected to affect interaction between histone H3 tail and the Oct4 acidic patch. To test this possibility, we assembled nucleosomes with H3K27ac-modified histone H3 and examined binding of Oct4 by native gel electrophoresis (**Fig. 3e, Extended Data Fig. 6a**). The H3K27ac modification showed only a small impact on binding of the first Oct4, which directly interacts with H3K27 residue, indicating that this interaction is not required for stability of the Oct4-nucleosome complex (**Fig. 3e**). However, binding of the second and third Oct4 was increased with H3K27ac nucleosomes compared to unmodified nucleosomes (**Fig. 3e**). Deacetylation of H3K27ac reduced binding of the second and third Oct4 to levels comparable with unmodified nucleosome (**Extended Data Fig. 6b, c**). Thus, cooperative Oct4 binding to OBS2/3 is increased by H3K27 acetylation.

This finding prompted us to examine whether the interactions of Oct4 with histone H3 contribute to DNA positioning by Oct4 in the Lin28B nucleosome. We first developed an assay to directly assess DNA positioning, taking advantage of an endogenous Mnl I restriction site between OBS1 and OBS2 (**Fig. 1b**); this site should be accessible to Mnl I when the Lin28B DNA is positioned as in our Oct4-bound nucleosome structure (**Fig. 3c**). Using Lin28B nucleosomes, we only observed partial digestion by Mnl I (**Fig. 3f, Extended Data Fig. 6d**), which is consistent with Lin28B DNA adopting multiple conformations on the nucleosome. In contrast, the Mnl I restriction site was fully accessible in Oct4-bound nucleosomes, indicating that Oct4 binding stabilizes the nucleosomal DNA in a conformation in which the Mnl I site is exposed (**Fig. 3f, Extended Data Fig. 6d**). H3K27ac did not alter the sensitivity of nucleosomes alone to Mnl I digestion (**Fig. 3f, Extended Data Fig. 6d**), however, Oct4-bound H3K27ac nucleosomes showed higher protection from Mnl I digestion compared to Oct4-bound unmodified nucleosomes (**Fig. 3f and Extended Data Fig. S6d**). These results suggest that Oct4 induces an inward movement of the DNA on the H3K27ac nucleosome, compared to unmodified nucleosome. Such movement might be induced by the loss of electrostatic interaction between the H3K27 residue and the Oct4 acidic patch, and it would likely have limited range, until Oct4 gets close to the nucleosome; further DNA movement would require DNA unwrapping or Oct4 dissociation. The DNA movement would increase exposure of OBS2/3 leading to higher binding of second and third Oct4 (**Fig. 3a**). To test this hypothesis, we digested Oct4-bound nucleosomes with MNase and sequenced the protected DNA (**Extended Data Fig. 6e**). We found that 70% of Oct4-bound unmodified nucleosomes were in a defined position, in agreement with our structural and biochemical data (**Fig. 3g**). In contrast, Oct4-bound H3K27ac nucleosomes were less well positioned, with a major species (20%) shifted by 1bp inwards (**Fig. 3g, Extended Data Fig. 6f**).

To mimic the changes in the DNA positioning caused by Oct4 on H3K27ac nucleosomes, we moved OBS2/OBS3 by 1 bp (Lin28B_OSO+1) or 2 bp (Lin28B_OSO+2) relative to OBS1. Modeling reveals that inward sliding of DNA for 1bp would expose binding site for Oct4_POU_S_ at both OBS2 and OBS3, thus changing interaction at OBS2 from Oct4_POU_HD_ to Oct4_POU_S_ (**Extended Data Fig. 6g**). We used unmodified nucleosomes bearing those constructs to test Oct4 binding and observed increased binding to OBS2/3 with Lin28B_OSO+1 construct, compared to Lin28B (**Extended Data Fig. 6h**). Lin28B_OSO+2 construct showed binding of Oct4 comparable to Lin28B (**Extended Data Fig. 6i**). These data show that Oct4 binds to Lin28B_OSO+1 in a manner similar to its binding to Lin28B on H3K27ac nucleosome, and support our conclusion that DNA movement of ~1 bp on Oct4-bound H3K27ac nucleosome increases binding to internal sites. To test this model further, we examined binding of Oct4 to H3K27ac Lin28B_OSO+1 nucleosomes, which would mimic a +2 bp movement, and observed binding comparable to Lin28B or Lin28B_OSO+2 nucleosomes (**Extended Data Fig. 6j**).

H3K27 methylation is a silent mark that would increase the bulkiness of the Lys residue, which could affect the interaction between histone H3 tail and the Oct4 acidic patch. To test if H3K27 methylation modulates Oct4 binding, we assembled nucleosomes with H3K27me3 and examined binding of Oct4 by native gel electrophoresis. We found that H3K27me3 did not significantly change Oct4 binding (**Extended Data Fig. 7a**). Consistent with that observation, we did not observe change in DNA positioning of Oct4 bound H3K27me3 nucleosomes by Mnl I digestion and MNase sequencing compared to unmodified nucleosomes (**Extended Data Fig. 6e, 7b, c**).

### H3K27ac increases Oct4:Sox2 cooperativity

In the Lin28B sequence, the Sox2-binding site is located between OBS2 and OBS3, forming a composite site with the latter. In our Oct4-bound nucleosome structure, the Sox2-binding site faces outwards, and modeling shows that Sox2 could bind to that site, with minor clashes with H2A (**Extended Data Fig. 7d**). Thus, Oct4 binding to the Lin28B nucleosome should facilitate Sox2 binding, by stabilizing the exposure of its binding site. We tested this hypothesis using native gel electrophoresis and observed that Sox2 could bind more efficiently to Oct4-bound Lin28B nucleosome than to Lin28B nucleosome alone (**Extended Data Fig. 7e-g**).

The inward DNA movement caused by Oct4 binding to the H3K27ac nucleosome would further increase exposure of the Sox2 site and alleviate the small clash between Sox2 and histone H2A (**Extended Data Fig. 7d**). Indeed, we found that Sox2 was able to bind better to Oct4-bound H3K27ac nucleosomes than to Oct4-bound unmodified nucleosomes (**Fig. 3h**). To validate our finding, we moved the Sox2-binding site to be 5bp closer to OBS2, which would reduce its exposure in Oct4-bound nucleosomes but not in unbound nucleosomes that can slide. We observed that this shifting of the Sox2-binding site strongly reduced binding of Sox2 to Oct4-bound H3K27ac nucleosomes, indicating that Oct4 binding to OBS1 determines DNA positioning and Oct4-Sox2 cooperativity (**Fig. 3i**). Notably, Sox2 binding to free H3K27ac nucleosomes was not affected when its binding site was shifted by 5b, showing that DNA slides on free nucleosome, transiently exposing the Sox2-binding site and allowing its binding. In contrast, when Oct is bound to the H3K27ac nucleosomes, DNA is positioned and cooperative binding with Sox2 is determined by the distance between OBS1 and the Sox2-binding site.

### Oct4 binding and cooperativity on other human sequences

To investigate whether our findings with the Lin28B nucleosome apply to other human DNA sequences, we assembled nucleosomes with a 186-bp-long DNA from the regulatory region near matrilin 1 gene ^5^ (nMatn1 DNA) (**Extended Data Fig. 8a**), which contains multiple Oct4 binding motifs. The initial cryo-EM reconstructions of the nMatn1 nucleosome in complex with Oct4 showed a density near the linker DNA, similar to the density of Oct4 bound to Lin28B DNA (**Extended Data Fig. 8b-g, Table S1**). Focused classification and refinements improved the resolution to 2.3 Å in the nucleosome portion (**Extended Data Fig. 8c-f**) and to 5.6 Å for a 25-kDa Oct4 region bound to the linker DNA (**Extended Data Fig. 8h-j**). We performed MNase sequencing to determine the position of the nMatn1 DNA on the Oct4-bound nucleosome and combined that information with the cryo-EM map to build a model for Oct4 bound to the nMatn1 nucleosome (**Extended Data Fig. 8k-m, 9a**).

Our structural and MNase sequencing data reveal that, despite the presence of multiple Oct4 motifs in the nMatn1 DNA, Oct4 predominantly binds one binding site in the linker DNA, near the nucleosome entry/exit site (mOBS1) (**Fig. 4a, Extended Data Fig. 9b**), a position overall similar to that in the Lin28B nucleosome structure. Both DNA-binding domains of Oct4 engage the nMatn1 nucleosome: while Oct4_POU_HD_ is bound close to the nucleosome dyad, Oct4_POU_S_ is located distally from the nucleosome (**Fig. 4a-b**). This arrangement differs from that with the Lin28B nucleosome, in which Oct4_POU_S_ was bound to the linker DNA, close to the nucleosome dyad, whereas Oct4_POU_HD_ was located distally from the nucleosome (**Fig. 1a, Extended Data Fig. 2h**). Nevertheless, despite its distal position relative to the nMatn1 nucleosome, Oct4_POU_S_ interacts with the histone H3 tail via a smaller acidic patch of Oct4 formed by side chains of helices 4 and 5 (**Fig. 4a-b, Extended Data Fig. 9c**). Notably, the H3 tail from the entry/exit site opposite to the Oct4-binding side interacts with Oct4 (**Fig. 4a-b, Extended Data Fig. 9c**), in contrast with the Lin28B nucleosome.

**Fig. 4.**
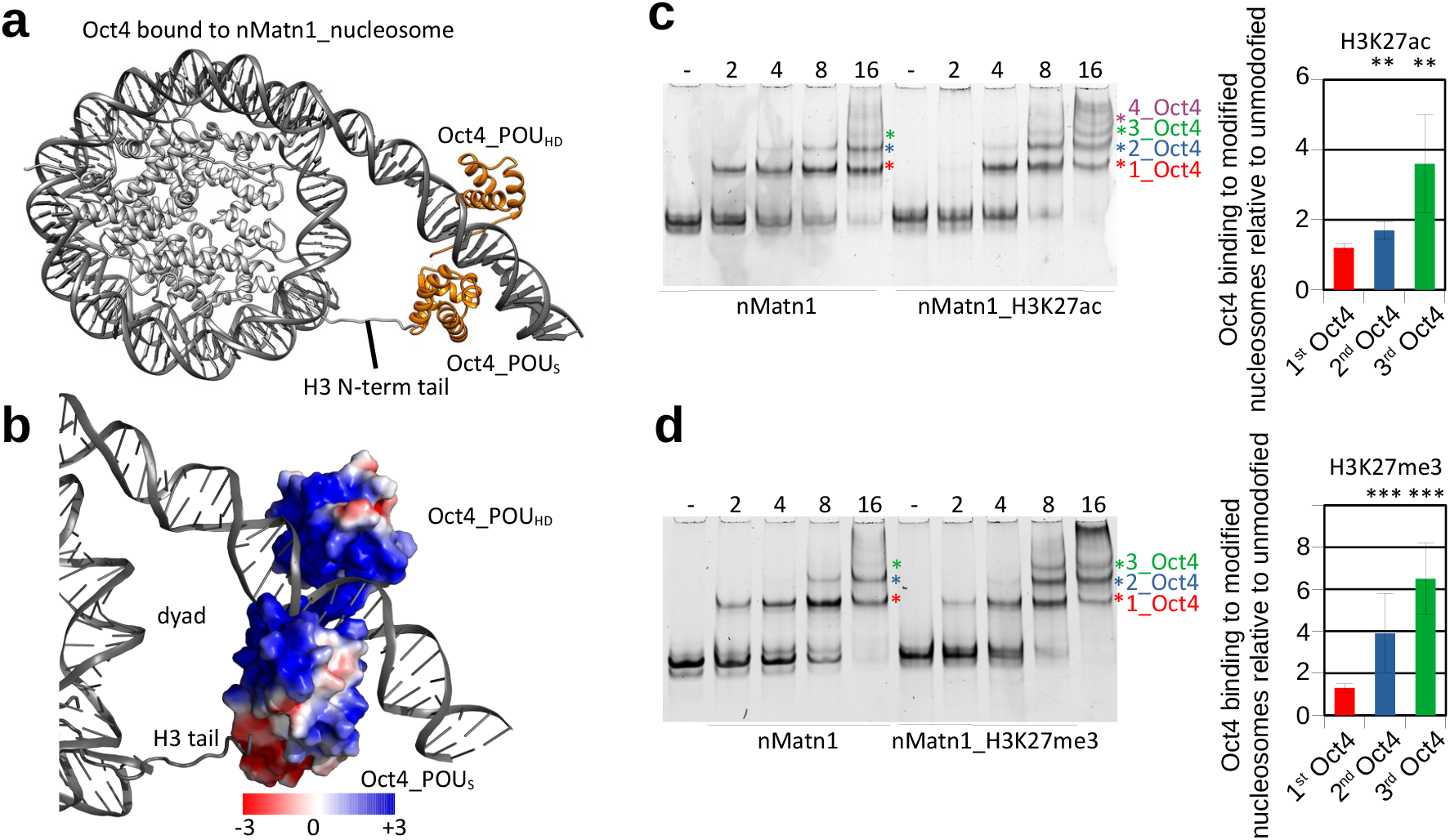
Histone modifications modulate Oct4 binding to nMatn1 nucleosome. **a)** A structural model of human Oct4 (orange) bound to a nucleosome (grey) assembled with a 186-bp DNA fragment from the nMatn1 regulatory element. **b)** Surface model showing electrostatic potential of Oct4_POU_S_ domain. The positively charged H3 tail is near a negatively charged surface of Oct4. **c)** Left, representative native gel electrophoresis showing Oct4 binding to the nMatn1 or nMatn1_H3K27ac nucleosomes. Colored asterisks indicate nucleosomes with different molecules bound: red, 1 Oct4; blue, 2 Oct4; green, 3 Oct4; purple, 4 Oct4. Right, quantification of the native gel electrophoresis, using bands marked with asterisks. Data shown as mean and s.e.m. of 4 independent experiments; ** p<0.01, Student’s t-test. **d)** Left, representative native gel electrophoresis showing Oct4 binding to unmodified or H3K27me3 nMat1n nucleosomes. Colored asterisks mark bands as in panel b. Right, quantification of the native gel electrophoresis, using bands marked with asterisks. Data shown as mean and s.e.m. of 4 independent experiments; *** p<0.001, Student’s t-test.

Oct4 interaction with the H3 tail in the nMatn1 nucleosome prompted us to test if H3K27 modifications also modulate histone cooperativity on this human sequence, as it does for the Lin28B sequence. We found that both H3K27ac and H3K27me3 modifications increased binding of the second and especially the third Oct4 (**Fig. 4c, d**).

Taken together, our biochemical and structural data reveal the mechanism for cooperativity of Oct4 and Sox2. Oct4 binding to OBS1 on Lin28B or nMatn1 nucleosomes stabilizes the positioning of nucleosomal DNA, to expose internal TF binding sites, thereby facilitating binding of additional Oct4 and of Sox2. The internal binding sites are distant from OBS1 (**Fig 3a, Extended Data Fig. 7d**), pointing to an allosteric mechanism for TF cooperative binding, mediated by DNA positioning on the nucleosome. Moreover, H3K27 modfications can modulate cooperativity of Oct4 and downstream factors, such as other Oct4 molecules or Sox2, by altering the interaction between the Oct4 acidic patch and the H3 tail, which in turn affects positioning of the nucleosomal DNA and exposure of internal binding sites for downstream factors.

## Discussion

Our data show that (1) Oct4 can directly modify nucleosomes by positioning DNA and by promoting their decompaction and (2) pioneer TF cooperativity can be regulated by histone modifications. These findings reveal how Oct4 can promote binding of downstream factors and suggest that the preexisting epigenetic landscape can tune TF activity to ensure proper cell reprogramming.

We were able to capture Oct4 bound to nucleosomes assembled with endogenous Lin28B and nMatn1 DNA by cryo-EM, which revealed how Oct4 binding causes nucleosomal DNA repositioning and exposure of internal TF binding sites and it also unveiled previously unknown Oct4 interactions with histones. A prior structure of Oct4 bound to engineered nucleosomes did not reveal interactions with histones ^21^, but those nucleosomes contained the strong 601 positioning sequence ^40^ with the Oct4-binding site inserted in a position that could have prevented interactions with H3 and H2A tails. Our data indicate that proper positioning of a TF’s DNA-binding site on the nucleosome is required for specific interactions and for formation of a stable Oct4-nucleosome complex. Our findings support a model in which initial binding of Oct4 to a partially exposed motif on the nucleosomal DNA leads to transient complexes that undergo DNA sliding to achieve stable Oct4 binding via its two DNA-binding domains (**Extended Data 10**). This model is consistent with recent *in vivo* data showing that pioneer TFs bind preferentially next to nucleosomes ^41^.

Our findings show that a pioneer TF can directly alter the chromatin environment by stabilizing DNA on the nucleosome. DNA sliding on the nucleosome can occur spontaneously or be facilitated by chromatin remodeling complexes. In fact, Oct4 and other TFs can recruit chromatin remodeling complexes ^42–46^, which might facilitate nucleosome sliding to properly position DNA binding motifs. Chromatin remodeler Brg1 is required for Oct4 binding to a subset of gene regulatory elements in cells ^13^, and inhibition of Brg1 catalytic activity reduces amount of already bound Oct4 at these elements *in vivo* ^25^, implying that chromatin remodelers support Oct4 by properly positioning nucleosome at those specific locations. Our findings suggest that at other sites, Oct4 binding itself can directly position nucleosomal DNA and alter the accessibility of sites for downstream factors.

Notably, we observe that Oct4 alters the conformation of histone H4 N-terminal tail, affecting inter-nucleosome interactions and promoting chromatin de-compaction. Recently, the Sox11 DNA-binding domain was proposed to affect the H4 tail position ^47^, but such mode of H4 regulation would limit TF binding to restricted regions on the nucleosome, where the DNA-binding domain would directly clash with the H4 tail. In contrast, our data reveal that the interaction between Oct4 and histone H4 tail involves the disordered activation domain of Oct4 and takes place 70 Å away from the site where its DNA-binding domains interact with the nucleosome, indicating that interaction with the H4 tail does not depend on the location of the Oct4 binding site. In agreement with our findings on Oct4, recent work suggests that activation domain of FOXA1 binds histones which is required for FOXA1 to open chromatin, although the mechanism of this interactions remain elusive ^48^.

Perhaps our most consequential finding is that TF binding and cooperativity can be regulated by histone modifications. Notably, our data show that H3K27 modifications did not affect binding of the first Oct4 to Lin28B or nMatn1 nucleosome, but it altered the cooperative binding of additional Oct4 or of Sox2 to nucleosomal internal sites. These findings are consistent with previous *in vivo* data correlating Oct4-binding sites with H3K27ac ^26, 27, 29^. However, positive correlation with histone marks has not been observed for FOXA2 and GATA4 ^27^, suggesting that not all TFs are able to recognize epigenetic marks. Our data reveal that histone modifications can modulate transcription factor binding and cooperativity. These findings suggest that preexisting epigenetic landscape can tune pioneer TF activity to ensure proper cell reprogramming.

## Methods

### Protein expression, mutagenesis and purification

The *Xenopus laevis* histones for nucleosome assembly were over-expressed in BL21(DE3) pLysS E. coli strain, and purified from inclusion body as described previously ^49^.

The cells were grown in LB medium at 37 °C and induced with 1 mM IPTG when OD (600) reached 0.6. After 3h of expression, the cells were pelleted down, resuspended in lysis buffer (50 mM Tris-HCl, pH 7.5, 150 mM NaCl, 1 mM EDTA, 1 mM DTT, 0.1 mM PMSF), and frozen. Later, the frozen cells were thawed and sonicated. The pellet containing inclusion bodies was recovered by centrifugation at 5000 rpm for 20 min at 4 °C. The inclusion body pellet was washed three times with lysis buffer containing 1% Triton X-100, followed by two washes with lysis buffer without Triton X-100.

Each histone protein was extracted from the purified inclusion body pellet in a buffer containing 50 mM Tris (pH 7.5), 2 M NaCl, 6 M guanidine hydrochloride, 1 mM DTT for overnight at room temperature. Any insoluble components were removed by centrifugation. Proteins making histone pairs (H2A and H2B, H3 and H4) were combined in equimolar ratios and dialyzed two times in 1 L of refolding buffer (25 mM HEPES/NaOH, pH 7.5, 2 M NaCl, 1 mM DTT) at 4 °C. Any precipitate was removed by centrifugation for 20 min, 13000 rpm at 4 °C. The soluble histone pairs were further purified via cation-exchange chromatography in batch (SP Sepharose Fast Flow resin). The samples were diluted four fold with buffer without salt (25 mM HEPES/NaOH, pH 7.5 and 1 mM DTT) and bound to the resin for 30 min. The resin was extensively washed with 500 mM salt buffer in batch (25 mM HEPES/NaOH, pH 7.5, 500 mM NaCl and 1 mM DTT) and loaded onto a disposable column. On the column, the resin was washed, and pure proteins were eluted with 25 mM HEPES/NaOH, pH 7.5, 2 M NaCl, 1 mM DTT. Soluble histone pairs were concentrated and purified on a Superdex S200 size exclusion column (GE) size exclusion column equilibrated in 25 mM HEPES/NaOH, pH 7.5, 2 M NaCl, 1 mM DTT. Clean protein fractions were pooled, concentrated and flash frozen.

For cryo-EM grid freezing of ‘assembly 1’ (see below), commercially available Oct4 from Abcam (<u>ab</u> <u>134876</u>) was used. The protein (Mr ~ 52 kD) is fused with the herpes simplex virus VP16 transactivation domain at the N-terminus and a 11R tag at the C-terminus. For the ‘assembly 2’ for cryo EM and all the other assays, His-tagged Oct4 (Mr ~ 39 kD) was expressed in a pET28 vector, and purified under denaturing conditions from inclusion body using Talon affinity resins. To refold the Oct4 protein, the first overnight dialysis was carried out in 2 M urea, 50 mM HEPES (pH 7.5), 250 mM NaCl, 50 mM L-Arginine, and 2 mM DTT. Then, the second and third dialysis was carried out for 1 h in a buffer containing 50 mM HEPES (pH 7.5), 100 mM NaCl and 1 mM DTT.

All the Oct4 variants were made using the inverse PCR strategy (iPCR). All the oligo primers used for mutagenesis were purchased from Integrated DNA Technology and are listed in the Table S2. The iPCR reactions were set up in a total volume of 25 μl. After amplification, 10 μl of purified PCR product was incubated with 5 U of T4 PNK in 20 μl of 1× T4 DNA ligase buffer for 1 h at 37°C. 200 U of T4 DNA ligase was added to the reaction and incubated for 1 h at room temperature. Finally, 10 U of Dpn I was added to the reaction and incubated for 1h at 37°C. From this mixture, 5 μl was used to transform the competent XL1-Blue *E. coli* cells. The clones were selected on Kanamycin plates, and were subsequently confirmed by sequencing.

### Histone octamer assembly and purification

Histone octamer purification was done using the standard protocol ^49, 50^. In brief, a 2.5-fold molar excess of H2A-H2B dimer was mixed with H3-H4 tetramer in the presence of buffer containing 2M NaCl (25 mM HEPES pH 7.5, 2M NaCl, 1 mM DTT). After overnight incubation at 4 °C, the assembled octamer was separated from excess dimer using a Superdex S200 Increase 10/300 GL column on an AKTA FPLC system. The fractions were analyzed on SDS-PAGE, pooled and concentrated for final nucleosome assembly.

### LIN28B 182bp DNA amplification

A custom synthesized (Integrated DNA Technology) 162 bp LIN28 genomic DNA ^6^ was cloned into pDuet plasmid. To make the longer 182 bp LIN28B DNA fragment by PCR, two primers were designed so that each contained an extra 10 base from the flanking genomic region of the canonical 162 bp LIN28 fragment used in previous studies ^6^.

The DNA sequence for the 182 bp extended DNA used in this study is:

5’: GCAT AAG TTA AGT GGT ATT AAC ATA TCC TCA GTG GTG AGT ATT AAC ATG GAA CTT ACT CCA ACA ATA CAG ATG CTG AAT AAA TGT AGT CTA AGT GAA GAA AGA AGG AAA GGT GGG AGC TGC CAT CAC TCA GAA TTG TCC AGC AGG GAT TGT GCA AGC TTG TGA ATA AAG ACA CAT ACT TCA T;3’

The underlined sequence shows the additional bases added to the 162 bp LIN28 fragment.

### Mutant LIN28B DNA

Custom synthesized 182 bp LIN28B DNA was purchased from IDT with following mutations in the three Oct4 binding sites:

LIN28B_1M (binding site 1 mutation): ATT AAC AT - GCGTCGAT

LIN28B_2M (binding site 2 mutation): ATT AAC AT - GCG GCT AT

LIN28B_3M (binding site 3 mutation): ATG CTG AAT - GCG GGT AA

The fragments were later PCR amplified to generate DNA for nucleosome assemblies.

### LIN28 DNA with shifted binding sites

OBS (+) 1 bp DNA:

GCAT AAG TTA AGT GGT ATT AAC ATA TCC TCA GTG GTG AGT **<u>A</u>** ATT AAC ATG GAA CTT ACT CCA ACAATA CAG ATG CTG AAT<u>AA</u> TGT AGT CTA AGT GAA GAA AGA AGG AAA GGT GGG AGC TGC CAT CAC TCAGAA TTG TCC AGC AGG GAT TGT GCA AGC TTG TGA ATA AAG ACA CAT ACT TCA T

OBS (+) 2 bp DNA:

GCAT AAG TTA AGT GGT ATT AAC ATA TCC TCA GTG GTG AGT **<u>AA</u>**ATT AAC ATG GAA CTT ACT CCA ACAATA CAG ATG CTG AAT<u>A</u> TGT AGT CTA AGT GAA GAA AGA AGG AAA GGT GGG AGC TGC CAT CAC TCAGAA TTG TCC AGC AGG GAT TGT GCA AGC TTG TGA ATA AAG ACA CAT ACT TCA T

SBS (-) 5 bp DNA

<u>GCAT AAG TTA</u> AGT GGT ATT AAC ATA TCC TCA GTG GTG AGT ATT AAC AT G GAA C TT A ACAATA **ACT CC** CAG ATG CTG AA T AAA T GT AGT CTA AGT GAA GAA AGA AGG AAA GGT GGG AGC TGC CAT CAC TCAGAA TTG TCC AGC AGG GAT TGT GCA AGC TTG TGA ATA AAG ACA <u>CAT</u> <u>ACT TCA</u> <u>T</u>

All the LIN28 variants with the binding sites shifted were purchased from IDT.

### Oct4 binding DNA sequences from human genome

The 186 bp nMTN (near matrilin 1 gene) sequence was selected from the human genome (https://www.ncbi.nlm.nih.gov/genome/gdv) from the position GRCh38:1:30216402:30217024:1 on chromosome 1 ^5^. The DNA fragment was selected based on the presence of the following Oct4 motifs: (ATGCTAAT, ATTAGCAT, ATTAACAT or ATGTTAAT).

The 186 bp nMatn1 sequence is shown below:

ACATGCACACATGCTAATATATGCACACAATGCACACAGGTTAATATATACACATACACACACATGCACACACACGTGCACACATATATGCACATGCATGCACACACGTATATGCACACACATGCACATGCATGCGCACATAGTCACACACATGCACACATTAGCATATGCATACACATACATGCA

The DNA was custom synthesized from IDT. The following primers were used for its PCR amplification in large scale: ACATGCACACATGCTAATATATG and TGCATGTATGTGTATGCATATGC.

### Nucleosome assembly

Nucleosome assembly was carried out using a ‘double bag’ dialysis method as described ^51, 52^. The histone Octamer and nucleosomal DNA fragment were mixed in equimolar ratios in a buffer containing 50 mM HEPES pH7.5, 2M NaCl, 2 mM DTT. The mixture was placed into a dialysis button made with a membrane of cut-off 3.5 kD. The dialysis button was placed inside a dialysis bag (6-8 kD cutoff membrane) filled with 50 ml of buffer containing 25 mM HEPES pH 7.5, 2M NaCl and 1 mM DTT. The dialysis bag was immersed into a 1L of buffer containing 25 mM HEPES pH 7.5, 1M NaCl, 1 mM DTT and dialyzed over-night at 4 °C. The next day, buffer was changed to 1L of a buffer with 25 mM HEPES pH 7.5 and 1 mM DTT, and dialysis was continued for 6-8 h. In the last step, the dialysis button was removed from the dialysis bag and dialyzed overnight into a fresh buffer without any salt (50 mM HEPES pH 7.5, 1 mM DTT). The nucleosome assemblies were assessed on a 6% native PAGE using SYBR Gold staining.

### Assembly of modified nucleosomes

H3K27ac nucleosomes were assembled using the LIN28B DNA (unlabeled or Cy5-labeled) and histone octamer with H3K27ac modification (custom purchased from Epicypher). For H3K27-trimethyl nucleosomes, H3K27C mutant histone was generated using site directed mutagenesis and later expressed and purified from E.coli. The H3K27C mutant histone thus obtained was tri-methylated using the MLA protocol ^53^ and purified using a PD10 column. This tri-methylated H3 was used with other histones for octamer assembly. The purified H3K27 trimethylated octamer was mixed with LIN28B and nMatn1 DNA for the assembly of H3K27me3 nucleosomes.

### Nucleosome array assembly

A 1022 bp genomic region from the LIN28 genomic site was synthesized by DNA synthesis (Codex, Protein Technology Center, St Jude Children’s Research Hospital). For nucleosome array reconstitution, the DNA fragment was amplified to a larger scale by PCR. For the assembly, the DNA and histone octamer were mixed in a 1:5 ratio.

Nucleosome array reconstitution was carried out using the ‘double bag dialysis’ salt dilution method described above (see “nucleosome assembly”).

The synthesized genomic DNA sequence used for Array assembly is following (LIN28 182 bp region shown in bold):

TATTATGATATTTAATGCATCTGTGGCTTAACACTTGTGAGAGTTACCAGCTTGAAAATGATGGTGTTGACTACCTCTTGAATCACATCTATCAACCACTGGCACCTACCACCAAGCTGGCTTCAATTAGTATGTGTTGCTTTTTGGTATTAACAACTAACCGTACTAGAGACCAAAGTGAACCCTGATTTTTATATGTCTTTAATAATGGTGTTTTATCTAGTGTTTTTAAATTATCCTGTGTAGTATTTAGATTACCTCATTGTCCATTTTGACTCATGTTGTTTACAAGTGAAAATAAAAACACTTGAACTGTATGTTTTTAAAAGACAAAAAAGGGGTAGATGTTTGGAATGCGTTTCACTCGCATGCAGTCATCTGGAGGGACTGAAGCACTGTTTGCCTTTCTGTACACTCTGGGTTTTATATTCTCATTTCATGCCTAATGTCTTATTCTGTCAATTATGGATATGTTGAGGTTTAAAAAAATTACTTGATTAAAAATAAAACATATAACGTTGGCATTTATGATGGGTTTTGGAGATTTTTTATGATGTGTGTTCTTACTAAGTTAAGTGAAAGGATAAAAGGCCATTTAGGCAGTGTTTTACATCAGTTGGA**GCATAAGTTAAGTGGTATTAACATATCCTCAGTGGTGAGTATTAACATGGAACTTACTCCAACAATACAGATGCTGAATAAATGTAGTCTAAGTGAAGGAAGAAGGAAAGGTGGGAGCTGCCATCACTCAGAATTGTCCAGCAGGGATTGTGCAAGCTTGTGAATAAAGACACATACTTCA**TGTAGTCAGAAGAGTGGTCTGAAGCAAAATATTCAAAGCCTATTGAAGAAATCATTTTAGAATTTTTCTACAAGTTTTTTCTAATAAAGAATTTGAAAAGGGCAAATTTTATCAGTACCAAATATCTAAAATACGAATGTAGATTTATCAAACCATGAATGCAGTATTGATGTACTGTAACAAACGATTTGGTAAAACTATTGAGTTCTTTAAATTGGCC

### Assembly of nucleosome-Oct4 complex for cryo-EM grid freezing

#### Lin28B complex

Equimolar mixture of histone octamer and LIN28B DNA (2 µM each) were mixed with 1 µM of Oct4 (abcam <u>ab134876)</u> and 3 µM of Sox2 (50 mM HEPES pH 7.5, 2M NaCl, 20% glycerol, 5 mM DTT). The assembly was carried out with four steps of buffer changes over 72 hrs. The buffer changes were carried out to dilute out the salt concentration from 2M starting concentration to a final solvent condition of no salt. The three buffers, used for the assembly dialysis, contained 50 mM HEPES pH 7.5, 2 mM DTT, and varying NaCl concentrations of 2M, 1 M and 0 respectively. After the assembly, the samples were centrifuged at 13000 rpm for 10 min at 4 °C to remove any precipitates. Following this, the sample was concentrated using a 10 kD centricon to the concentrations needed for cryo EM grid freezing (0.5-1 µg/µl).

The assemblies were checked on 6% native gels followed by native Western blot analysis (**Fig. S1A**). For detection of nucleosome, Oct4 and Sox2, anti-H3, anti-Oct4 and anti-His antibodies were respectively used (see the Western blot detection section below).

#### nMatn1 complex

For the Oct4 bound to nMatn1 nucleosome, 1 µM of pre-assembled nucleosomes were mixed with 2 µM of His-tagged Oct4 (see above) and incubated at room temperature for 30 min. The sample was then transferred to ice until grid freezing.

### Restriction enzyme Mnl I digestion assays

For digestion of LIN28B nucleosomes, different dilutions of Mnl I (NEB) were made in the 1x CutSmart buffer (NEB). The digestion was carried out for 30 min at 25 °C. For the experiments involving Oct4 and Oct4 variants, the protein was incubated with nucleosome at 25 °C for 5 min before Mnl I addition. After the MnlI addition, the sample were kept at 25 °C for 30 min. After the digestion, the samples were run on a 6 % polyacrylamide gel to separate all the products, and then imaged by SYBR gold staining on a Typhoon scanner (**Fig. S5A**).

### Magnesium precipitation assay

LIN28B Nucleosome samples were incubated in varying MgCl_2_ concentrations for 10 min at 25 °C. The precipitated nucleosomes were separated from soluble nucleosomes by spinning at 10000 rpm for 10 min at 25 °C. The same procedure was followed for nucleosome samples containing WT Oct4 and other variants. However, for the experiments done in the presence of Oct4 and Oct4 variants, the nucleosomes were first mixed with 5-fold molar excess of Oct4 (or Oct variants) and kept at 25 °C for 5 min before any MgCl_2_ addition.

### Binding assays

The binding assays with Oct4 were performed at 25 °C in 50 mM HEPES (pH 7.5), 200 mM KCl, 1 mM DTT, 0.005% NP40. The binding assays involving both Oct4 and Sox2 were performed in 50 mM HEPES (pH 7.5), 1 mM DTT. Typically, 20-40 nM of nucleosome was incubated with different amounts of proteins (Oct4, Oct4 variants and Sox2). For the Oct4 binding experiments, the reaction was incubated for 10 min. For binding involving both Oct4 and Sox2, the reaction was incubated for 10 min after Oct4 addition, following which Sox2 was added and kept for an additional 5 min. The bound and unbound species were separated on a 5% or 6 % native polyacrylamide gel and imaged for Cy5 fluorescence using a Typhoon scanner. In experiments with nucleosomes without Cy5 label, SYBR gold staining was used to visualize the gels.

### Analysis of gels

All the gels were analyzed using Quantity One Basic version (Bio-rad). The data were exported and analyzed/plotted using Igor Pro (WaveMatrix Inc.). All the bands were selected using boxes of same size. The background correction was done separately for bands from each lane using boxes of identical size in the same lane.

#### Analysis of Mnl I digestion of nucleosomes

In case of nucleosome only experiment, after background correction the signal from the nucleosome band from each concentration point was normalized to the signal from the nucleosome lane in 0 Mnl I lane. For Mnl I digestion in presence of Oct4 or its variants, the signal of the Oct4 bound band from each of the Mnl I concentration was background corrected and then normalized to the signal of Oct4 bound band from the 0 Mnl I lane.

#### Analysis of Mg^2+^ precipitation assays

The relative compaction was calculated as the fraction of precipitated nucleosomes. For this, the following formula was used: Relative compaction = S_0_ – S_obs._ S_0_ is the signal of nucleosome band in 0 Mg^2+^ concentration normalized to 1, S_obs_ is the signal of all the soluble nucleosome bands normalized to signal of nucleosomes in 0 Mg^2+^ concentration. For precipitation experiments in presence of Oct4 or its variants, the signals from both the bound and unbound nucleosomal species were summed to calculate the soluble nucleosomes.

#### Analysis of Oct4 binding to wt LIN28B vs LIN28B_1M mutant

For binding to wt LIN28B nucleosomes we used the following equation: 2^nd^ Oct4 = [2_Oct4] / ([1_Oct4]+[2_Oct4]), where 2_Oct4 represents nucleosome with 2 Oct4 bound, and 1_Oct4 is a nucleosome with 1 Oct4 bound. [1_Oct4]+[2_Oct4] represent input Oct4 bound nucleosomes, which are substrates for binding of the second Oct4. For binding to LIN28B_1M nucleosomes we used the following equation: 2^nd^ Oct4 = [1_Oct4] / [Nucleosome], where Nucleosome represents input nucleosomes.

#### Analysis of Sox2 binding to wt LIN28B vs LIN28B_1M mutant

Binding of Sox2 to Oct bound nucleosome in Fig. is calculated as fraction of Sox2 bound to Oct bound LIN28B nucleosome: Sox = [1_Oct4_Sox2] / ([Oct4]+[1_Oct4_Sox2]); where 1_Oct4_Sox2 represent nucleosomes with both Oct4 and Sox2 bound, and Oct4 represents Oct4 bound nucleosomes. ([1_Oct4]+[1_Oct4_Sox2]) represents input Oct4 bound nucleosomes, which are substrates for binding of the Sox2 to Oct4 bound nucleosomes. Binding of Sox2 to Lin28B nucleosome is shown as a fraction of free Lin28B nucleosomes: Sox2 = [Sox2] / ([Nucleosome]-[1_Oct4]-[1_Oct4_Sox2]); where Sox2 represents Sox2 bound nucleosomes, Oct4 represents Oct4 bound nucleosomes, and Nucleosome input nucleosomes. Fraction of Sox2 bound to LIN28_1M mutant nucleosome is calculated as: Sox2 = [Sox2] / ([Nucleosome]-[1_Oct4])

#### Analysis of Oct4 binding to unmodified, H3K27ac and H3K27me3 nucleosomes

For binding to modified nucleosomes we used the following equation: 1^st^ Oct4 = ([1_Oct4]+[2_Oct4]+[3_Oct4]) / input nucleosome; 2^nd^ Oct4 =([2_Oct4]+[3_Oct4]) / 1^st^ Oct4 ([1_Oct4]+[2_Oct4]+[3_Oct4]); 3^rd^ Oct4 = [3_Oct4] / 2^nd^ Oct4 ([2_Oct4]+[3_Oct4]), where 1_Oct4 represents nucleosome with 1 Oct4 bound, 2_Oct4 is a nucleosome with 2 Oct4 bound, and 3_Oct4 is nucleosome with 3 Oct4 bound. Binding of 1^st^ Oct4 is shown relative to input nucleosome as substarte, binding of 2^nd^ Oct4 is show relative to 1^st^ Oct4 (substrate), and binding of 3^rd^ Oct4 is shown relative to 2^nd^ Oct (substrate). The quantification is shown as ratio of modified nucleosomes vs unmodified.

#### Analysis of Sox2 binding to H3K27ac nucleosomes

For binding to modified nucleosomes we used the following equation: Sox2 = [1_Oct4_Sox2] / [1_Oct4_Sox2] + [1_Oct4]; where 1_Oct4 represents nucleosome with 1 Oct4 bound, 1_Oct4_Sox2 is a nucleosome with Oct4 and Sox2 bound. Binding of Sox2 to Oct4 bound nucleosomes is shown relative to 1^st^ Oct4 bound nucleosomes (substrate). The quantification is shown as ratio of modified nucleosomes vs unmodified.

### Western blot detection

SDS-PAGE gels or native PAGE gels were transferred to PVDF membrane and blocked in TBST (50 mM Tris/HCl, pH 7.5, 150 mM NaCl, 0.1% tween-20) containing 5% milk for 1 h. Membranes were then incubated in primary antibody in TBST containing 5% milk for 1 h at room temperature. The membranes were washed 3 x for 5 min with TBST and incubated in secondary antibody for 1 h at room temperature. Membranes were washed for 3 x (~5 min each) with TBST before chemiluminescent detection. The following antibodies were used: anti-Oct4 antibody (Abcam), HRP-conjugated anti-His antibody (Invitrogen – Thermo Fisher), anti-H3 antibody (abcam ab1791) and anti-Sox2 antibody (Abcam ab 92494), HRP-conjugated anti-rabbit secondary antibody (Biorad).

### MNase-seq

Oct4 was bound to unmodified, H3K27ac or H3K27me3 nucleosome with Lin28B or nMatn1 DNA (20 mM HEPES pH 7.5, 50 mM KCl, 2.5 mM MgCl_2_ and 5 mM CaCl_2_) and digested by MNase (NEB) for 5 min at 25°C. MNase digestion was terminated by 50 mM EDTA. Cleaved nucleosome was subjected to phenol/chloroform extraction followed by ethanol precipitation of nuclesomal DNA and used for library preparation. Sequencing library was prepared using NEBNext® Ultra™ II DNA Library Prep Kit following manufacture’s manual. Amplification of library for illumine sequencing was performed by PCR using NEBNext® Multiplex Oligos for Illumina kit. Sequencing was pair end with 100 bp length. Paired reads were merged and filtered by the length of reads between 144bp and 146bp and mapped to LIN28B or nMatn1 sequence with Qiagen CLC genomics Workbench 20 software.

### MiDAC purification

MiDAC was purified from 1.25 L of adherent Flp-In™ 293 T-REx cell lines stably transformed with FlpIn^TM^ expression vector carrying FLAG-ELMSAN1/MIDEAS. The cells were grown in DMEM media (Gibco) supplemented with 10% FBS, 100 µg/ml hygromycin and induced for 24 hrs with 1µg/ml doxycycline (Thermofischer scientific). Cells were harvested and lysed using classical Dignam protocol ^54^ (doi:10.1101/pdb.prot5330). The complex was isolated from the nuclear fraction using anti-FLAG M2 beads from Sigma aldrich. The nuclear fraction was mixed with washed FLAG M2 beads and incubated overnight at 4 °C. Next day, the beads were washed with wash buffer (20 mM HEPES pH 7.9, 300mM NaCl, 1.5 mM MgCl2, 10% glycerol, 0.5 mM DTT, Protease inhibitors (Sigma)) four times. The complex was eluted from the beads in the elution buffer (20 mM HEPES pH 7.9, 100 mM NaCl, 1.5 mM MgCl2, 0.5 mM DTT, Protease inhibitors (Sigma)) after 30 minutes incubation at 4°C. This complex was flash frozen in liquid nitrogen and stored at −80°C.

### Deacetylation of H3K27ac nucleosomes

H3K27ac nucleosomes were deacetylated by human MiDAC deacetylase complex. The deacetylation reaction was carried out for 18 h at 25 °C in the following buffer: 50 mM HEPES (pH7.5), 100 mM KCl, 0.2 mg/ml BSA. A control parallel reaction containing H3K27ac nucleosomes, but no MiDAC, was also carried out under the identical conditions. The extent of deacetylation was confirmed by Western blot using anti-H3K27ac antibody.

### Negative stain electron microscopy

For the experiment looking at array compaction, 20nM of the LIN28 array was mixed with MgCl_2_ to a final [Mg^2+^] of 3 mM. For analysis of the effect of WT and ΔN Oct4 proteins, 70 nM (WT) and 100 nM (ΔN) proteins were respectively used with the mixture of array and MgCl_2_.

After ~ 10-15 min incubation at 25 °C, 3 µl of the sample was added to Lassey Carbon or quantifoil grids for 1 min, blotted dry and stained. For staining, 4 separate drops (~ 40 µl) of Uranyl Acetate / Uranyl Formate were added to a parafilm strip. The grid was briefly brought into contact with the stain for the first three drops before quick blotting. The last drop of stain was kept in contact with the grid for 1 min before the final blot drying.

The dried grids were imaged on a Talos L 120C microscope (Thermo Fisher Scientific) at the cryo-EM facility at St Jude Children’s Research Hospital. Several images were acquired at 73k - 92k magnification from regions showing good particle distribution. Specifically, a magnification of 73k was used for experiments involving Mg2+ compacted arrays in absence and presence of Oct4; for experiment with ΔN variant of Oct4 a magnification of 92k was used. The pixel size was 1.94 Å (73k) - 1.54 Å (92k) per pixel on the object scale. The images were later analyzed using the Image J software after matching the scale from the EM images.

### Negative stain Image analysis

Several particles were picked using Relion (N=450 for arrays in 3 mM MgCl_2_, N = 262 for array in 3mM MgCl_2_ with Oct4, and N = 307 for arrays in 3 mM MgCl_2_ with ΔN variant of Oct4). For particle picking, the images from the microscope were binned two-fold in Relion, and saved as 400 pixel x 400 pixel tiff images. The tiff images of the picked particles were later analyzed using the Image J software ^55^. First, the particles were encircled using the free-form selection tool in Image J. Later, the ‘set scale’ tool in Image J was used to set the size of the pixel in the image to 0.4 nm (0.2 nm pixel size at 73k magnification multiplied by 2 for binning in Relion). The particle sizes were measured using the image analyzer option in Image J and plotted.

### CryoEM grid preparation and data collection

For cryo-EM of Oct4 bound Lin28B nucleosome structure we have assembled Oct4_Sox2_Nucleosome complex as described. The sample was concentrated to 0.25 mg/ml for cryo-EM grid. To avoid the extensive aggregation of the complex sample on the cryo-EM grid, Oct4 and Sox2 were mixed with nucleosomes in 0.5:1 ratio during the assembly. Oct4 bound nMatn1 nucleosome was assembled as described with 2:1 ratio of Oct4 to nucleosome. 3 μl of the complex sample were applied to freshly glow-discharged Quantifoil R2/1 holey carbon grid. Humidity in the chamber was kept at 95% and temperature at + 10 °C. After 5 s blotting time, grids were plunge-frozen in the liquid ethane using FEI Vitrobot automatic plunge freezer.

For Lin28B nucleosome and Oct4 bound Lin28B nucleosome electron micrographs were recorded on FEI Titan Krios at 300 kV with a Gatan Summit K3 electron detector (~6000 and ~11000 micrographs) at the Cryo-EM facility at St. Jude Childrens’s Research Hospital. Image pixel size was 1.06 Å per pixel on the object scale. Data were collected in a defocus range of 7 000 Å – 30 000 Å with a total exposure of 90 e/Å^2^. 50 frames were collected and aligned with the MotionCorr2 software using a dose filter ^56, 57^. The contrast transfer function parameters were determined using CTFFIND4 ^58^. For Oct4 bound nMatn1 nucleosome, the data were recorder on FEI Titan Krios at 300 kV with a Falcon 4 electron detector (~35 000 micrographs) at the Cryo-EM facility at Dubochet Center for Imaging (DCI) at EPFL and UNIL. Data were collected in a defocus range of 7 000 Å – 25 000 Å. Image pixel size was 0.83 Å per pixel on the object scale.

Several thousand particles were manually picked and used for training and automatic particle picking in Cryolo ^59^. Particles were windowed and 2D class averages were generated with the Relion software package ^60^. Inconsistent class averages were removed from further data analysis. The initial reference was filtered to 40 Å in Relion. C1 symmetry was applied during refinements for all classes. Particles were split into 2 datasets and refined independently and the resolution was determined using the 0.143 cut-off (Relion auto refine option). All maps were filtered to resolution using Relion with a B-factor determined by Relion.

Initial 3D refinement was done with 2 600 000 particles. To improve the resolution of this flexible assembly, we have used focused classification followed by focused local search refinements. Nucleosome was refined to 2.8 Å. Density modification in Phenix improved the map to 2.5 Å ^61^. Oct4 bound to DNA (30 kDa) was refined to 4.2 Å using a subset of 65 000 particles after extensive sorting. Using density modification in Phenix, we have improved resolution and appearance of this density to 3.9 Å. The maps have extensive overlapping densities that we used to assemble the composite map and model. Lin28B nucleosome sample contained 1 000 000 particles, which were refined to 3.1 Å, and improved with density modification to 2.8 Å.

For the second dataset we collected 1400 images, yielding 68 000 nucleosomal particles, which refined to 3.7 Å. Classification revealed that approximately 21 000 particles has Oct4 bound, which refined to 4.2 Å.

Molecular models were built using Coot ^62^. The model of the NCP (PDB:6WZ5) ^63^ was refined into the cryo-EM map in PHENIX ^64^. The model of the Oct4 bound to DNA (PDB:3L1P) ^19^ were rigid-body placed using PHENIX, manually adjusted and re-build in COOT and refined in Phenix. Visualization of all cryo-EM maps was done with Chimera ^65^.

## Acknowledgments

We thank Cryo-EM facility members at St. Jude Children’s Research Hospital for support with grid screening, especially Alexander Myasnikov and Liang Tang for support with data collection of Lin28B nucleosome and Oct4 bound Lin28B nucleosome. We thank Alexander Myasnikov and Bertrand Beckert from Dubochet Center for Imaging (DCI) at EPFL and UNIL for data collection of Oct4 bound to nMatn1 nucleosome. We thank Inês Chen for critical reading and comments. Work in the Halic laboratory is funded by St. Jude Children’s Research Hospital, the American Lebanese Syrian Associated Charities and NIH awards 1R01GM135599-01 and 1R01GM141694-01.

## Author contributions

K.S. and M.H. designed the experiments. K.S. performed biochemical experiments and electron microscopy. S.B built models. J.D. performed MNase-seq experiments. D.M. purified MiDAC complex. K.S, S.B, J.D. and M.H. analyzed the data. K.S., S.B and M.H. wrote the paper.

## Data availability

EM densities have been deposited in the Electron Microscopy Data Bank under accession codes EMD-xxx, EMD-xxx, EMD-xxx, EMD-xxx, EMD-xxx, EMD-xxx and EMD-xxx. The coordinates of EM-based models have been deposited in the Protein Data Bank under accession codes PDB xxx. The RAW data are provided as a Supplementary Fig. file. All other data are available from the corresponding author upon reasonable request.

## Supplemental Information

**Extended Data 1.**
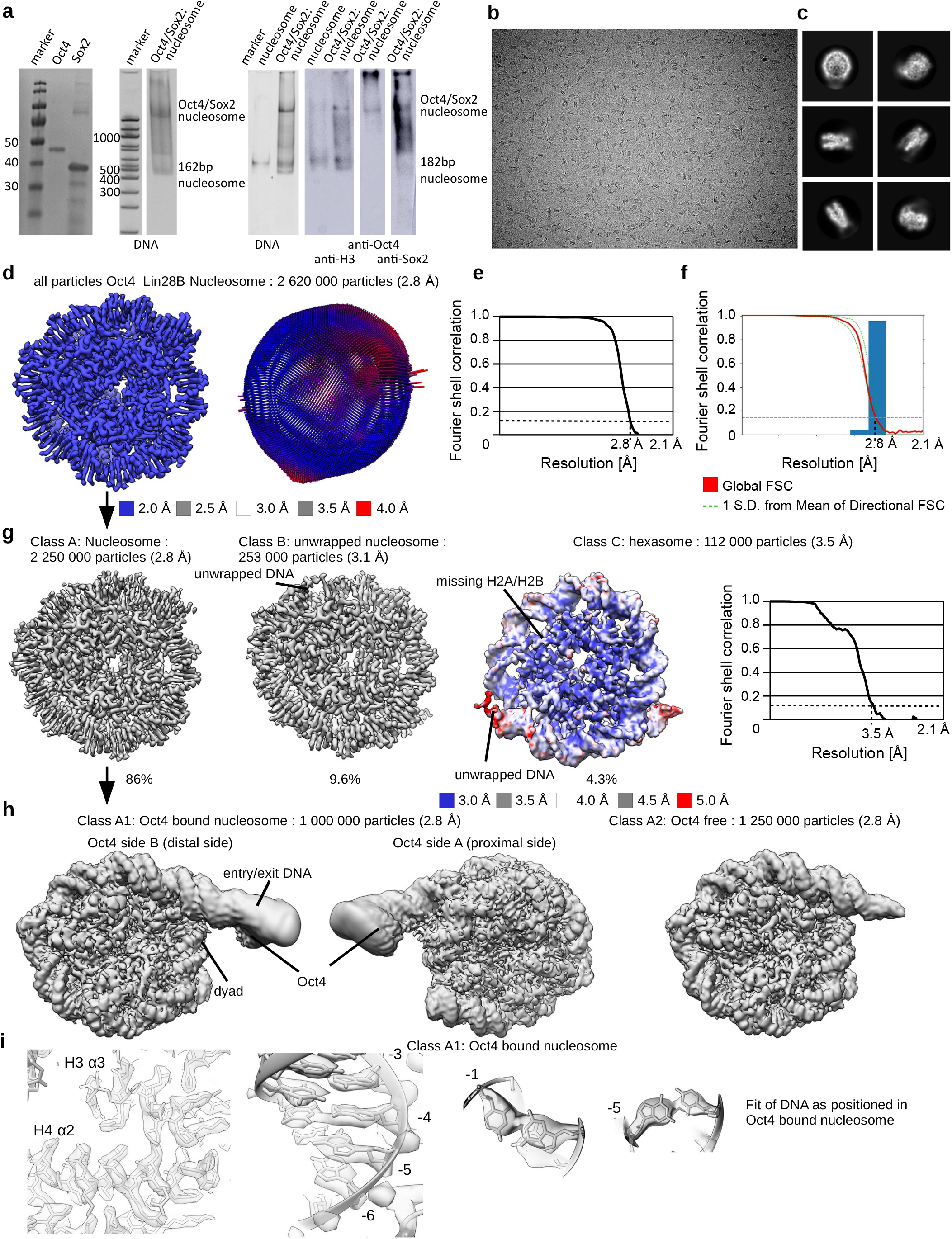
Assembly and cryo-EM of Oct4 bound to Lin28B nucleosomes. **a)** SDS-PAGE showing purification of Oct4 and Sox2 and the assembly of the Oct4_Sox2_nucleosome complex. From left: A SDS gel showing purification of Oct4 and Sox2 used in the experiments; a native gel stained for DNA showing the assembly of the Oct4_Sox2_nucleosome complex; western blots with anti-H3 antibody, anti-Oct4 antibody and anti-His antibody (Sox2). **b)** Representative cryo-EM micrograph collected with Titan Krios electron microscope at 300 keV. Nucleosome particles in multiple orientations are visible. **c)** Representative 2D class averages showing nucleosomes. Many details in nucleosomes are visible in 2D class averages. **d)** Cryo-EM map of nucleosome from the entire dataset, refined to 2.8 Å. The map is colored by local resolution. The model of the NCP (PDB:6WZ5) was refined into the cryo-EM map. Angular distribution for nucleosome is shown on the right. **e)** Fourier shell correlation (FSC) curve showing the resolution of the map in d). **f)** Directional FSC plot showing uniform resolution in all directions. **g)** Classification of the data in b) resulted in three major classes of nucleosomes and nucleosome like particles (nucleosome, unwrapped nucleosome and hexasome). Hexasome map is colored by local resolution and the FSC curve is shown on the right. Number of particles corresponding to each class is indicated. **h)** Classification of the nucleosome subset from g). Classification revealed two classes, nucleosome and nucleosome with bound Oct4. We did not observe density for Sox2. **i)** Left: the representative region showing map quality and fit of the model is shown for the nucleosome with bound Oct4. Right: bases in the DNA are well resolved.

**Extended Data 2.**
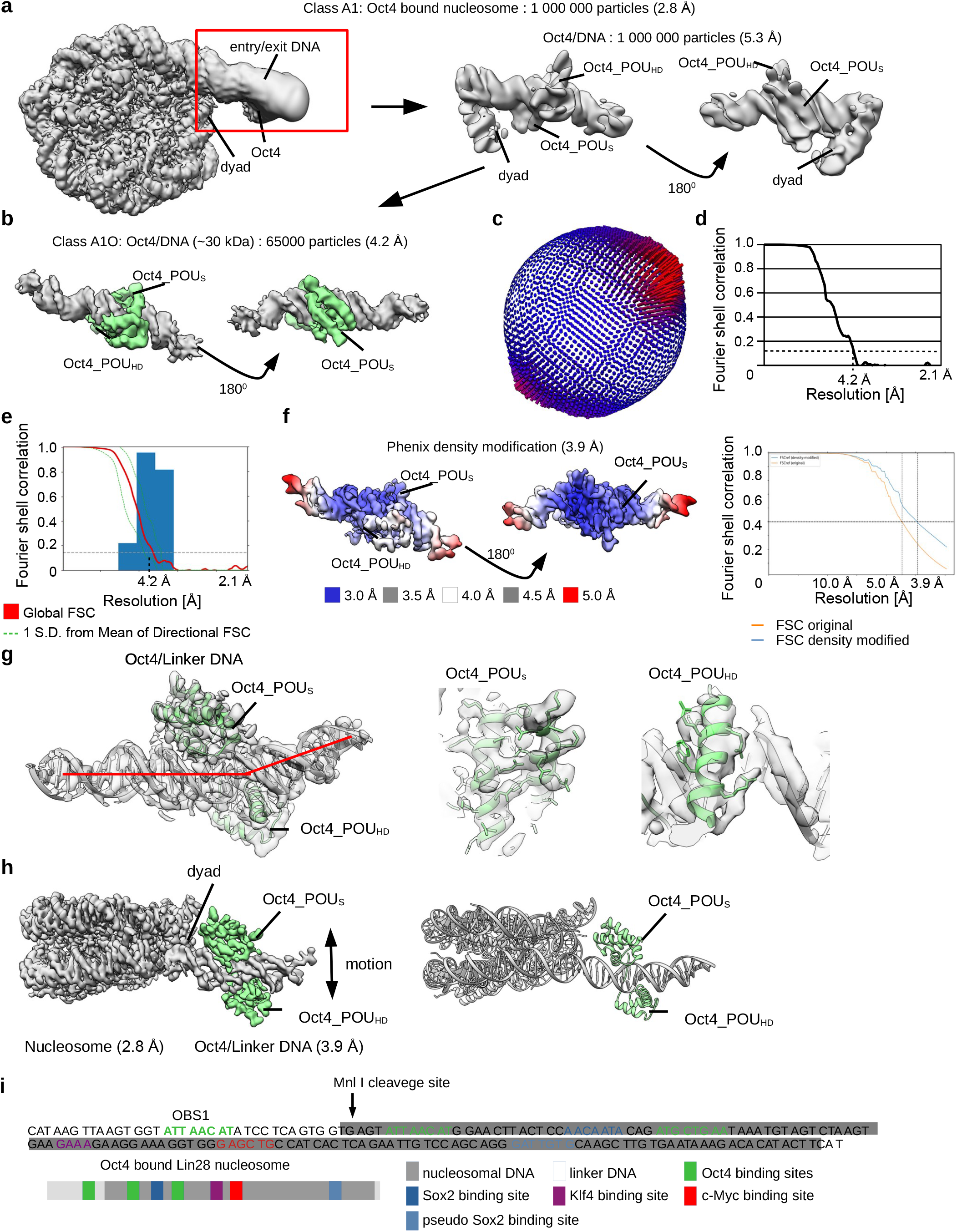
Classification of the Oct4_nucleosome complex. **a)** Focused refinement of the Oct4 density from the Oct4_nucleosome (left) complex improved the resolution to 5.3 Å (right). **b)** Cryo-EM map of Oct4 region from the Oct4_nucleosome complex. Focused classification and refinements improved the resolution of this 30 kDa fragment to 4.2 Å. **c)** Angular distribution for Oct4. **d)** The fourier shell correlation (FSC) curve showing the resolution of the map. **e)** Directional FSC plot showing uniform resolution in all directions. **f)** The Oct4_DNA density from b) was modified in Phenix, which improved the resolution to 3.9 Å. The map is colored by local resolution. Fourier shell correlation (FSC) curve showing the resolution is shown on the right. **g)** The model of the Oct4 bound to DNA (PDB:3L1P) was refined into the cryo-EM map. The representative region showing map quality and fit of the model is shown on the right. Red line shows the kink in the DNA. **h)** A composite cryo-EM map of Oct4 bound to the Lin28B nucleosome containing 182bp of DNA at 2.8-3.9 Å resolution (left). Model for the cryo-EM structure is shown on the right. **i)** DNA sequence and schematic representation showing Lin28B DNA positioning on the Oct4_nucleosome complex. Oct4, Sox2, Klf4 and c-Myc binding sites are labeled. The cleavage site for the restriction enzyme Mnl I (Fig. 3c) is marked with an arrow.

**Extended Data 3.**
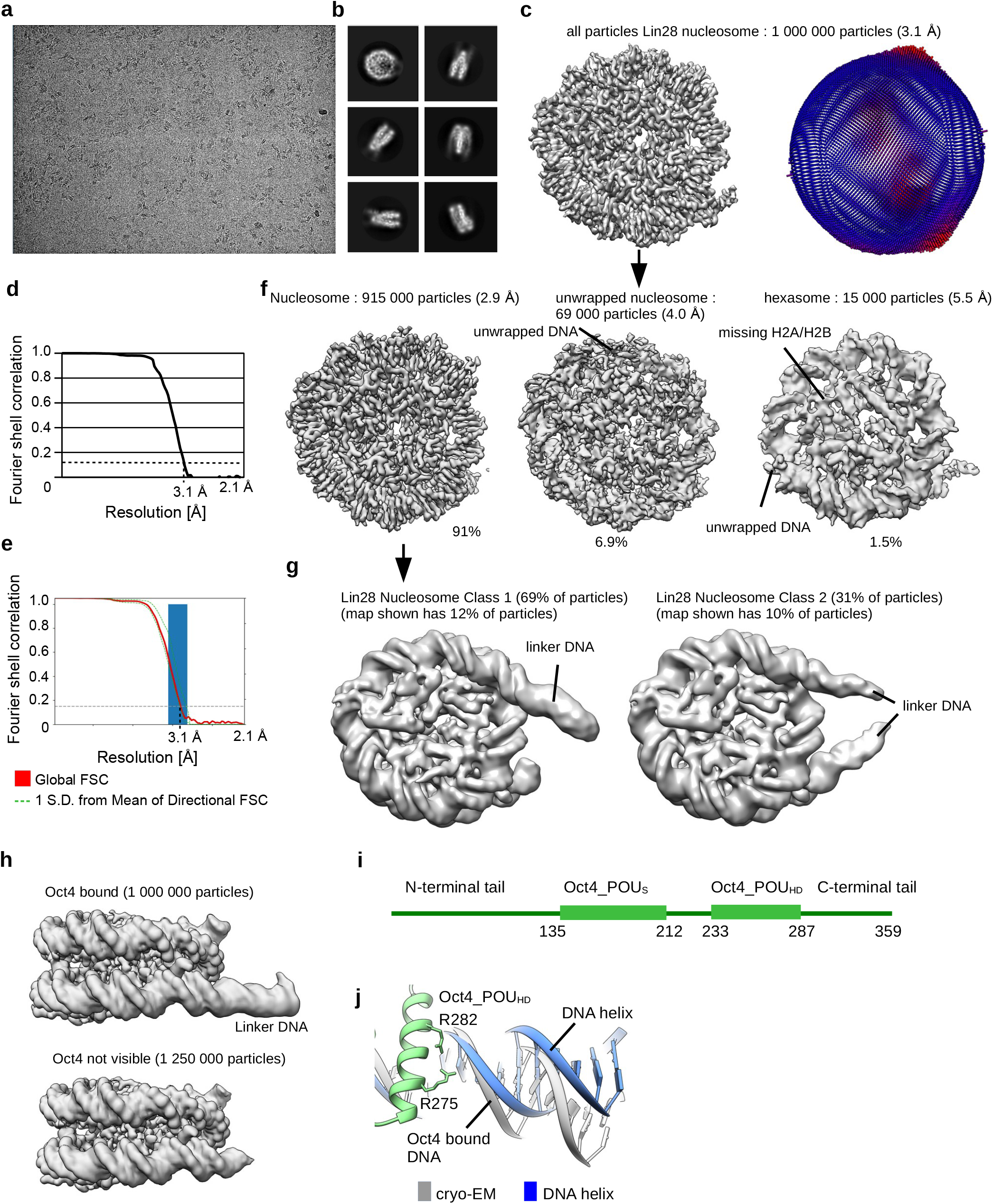
Cryo-EM of Lin28B nucleosome. **a)** Representative cryo-EM micrograph collected with Titan Krios electron microscope at 300 keV. Nucleosome particles in multiple orientations are visible. **b)** Representative 2D class averages showing nucleosomes. Many details in nucleosomes are visible in 2D class averages. **c)** Cryo-EM map of nucleosome from the entire dataset, refined to 3.1 Å. Angular distribution for nucleosome is shown on the right. **d)** Fourier shell correlation (FSC) curve showing the resolution of the map in c). **e)** Directional FSC plot showing uniform resolution in all directions. **f)** Classification of the Lin28B nucleosome dataset resulted in three classes of nucleosome like particles (nucleosome, unwrapped nucleosome and hexasome). Number of particles corresponding to each class is indicated. **g)** Classification of Lin28B nucleosome particles from f) showing that DNA protrudes on both sides of nucleosome in some classes. Protruding DNA gives prominent signal and can be reliably classified. **h)** Cryo-EM maps of Oct4_nucleosome complex and Lin28B nucleosome. DNA extends only on Oct4 bound side of the nucleosome in the Oct4 bound sample. Linker DNA is more defined and stabilized by Oct4. **i)** Schematic representation of Oct4 domains. The POU_S_ and POU_HD_ domains are structured, whereas the N- and C-terminal tails are disordered. **j)** Model of Oct4 bound to the linker DNA showing the kink in the DNA introduced by binding of Oct4_POU_HD_. Arg in Oct4_POU_HD_ interact with DNA.

**Extended Data 4.**
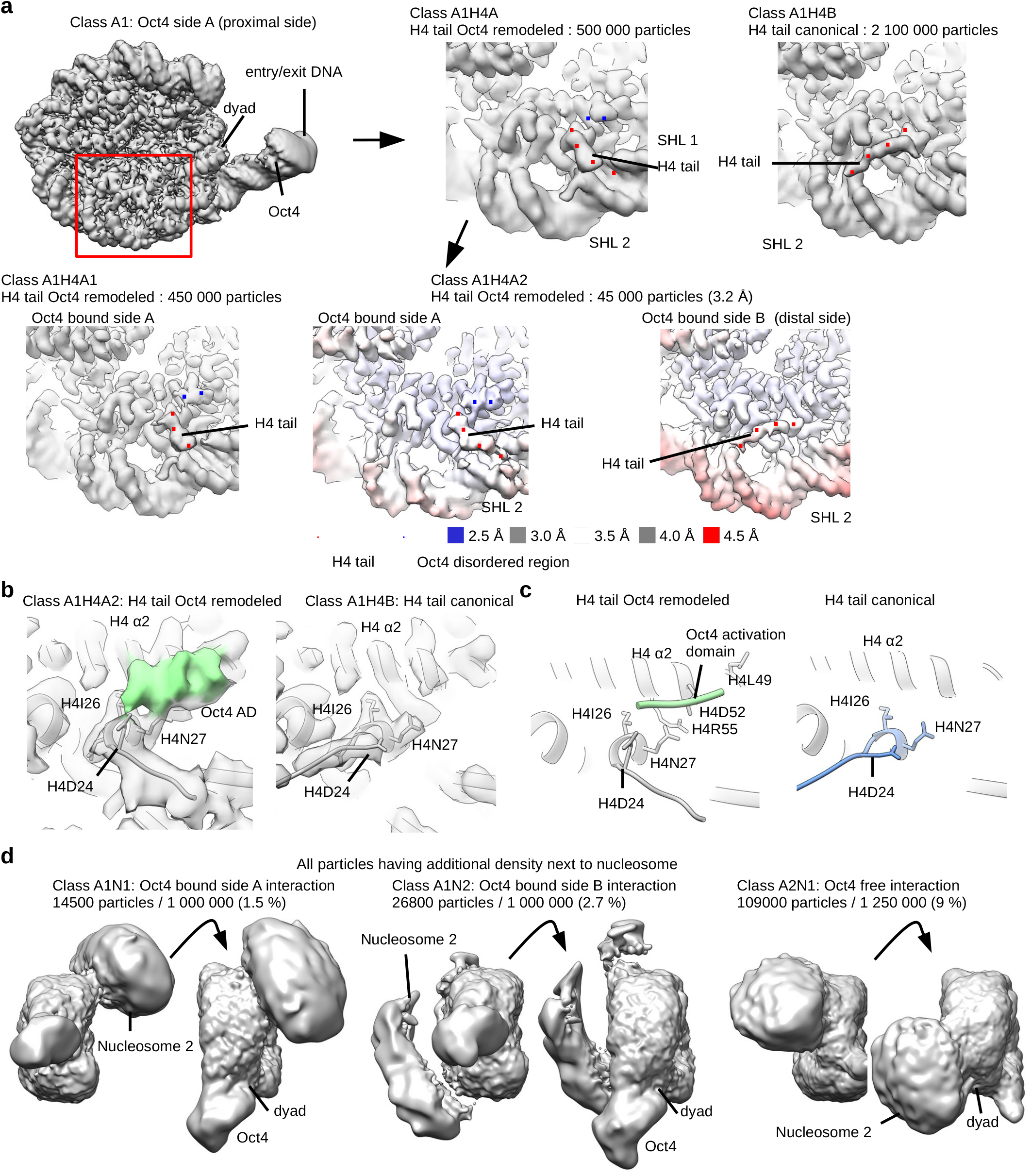
Intrinsically disordered region of Oct4 binds to histone H4. **a)** Classification of the cryo-EM data revealed two conformations of the H4 tail on the Oct4 proximal side. Red dots depict the H4 tail. The class with the re-positioned H4 tail, which goes to the SHL1, contains an additional density which is labeled with blue dots. Maps are colored by local resolution. **b)** Cryo-EM maps and fitted models showing two positions of the H4 tail. Note different orientation of H4D24 which interacts with the Oct4 density. Oct4 density is shown in green. **c)** Detailed views of the H4 α2 and N-terminal tail on the Oct4-proximal side of the nucleosome, showing the two distinct conformations of the H4 tail, canonical (blue, left) and Oct4 remodeled (grey, right). The disordered region of Oct4 that interacts with H4 is represented in green, and it contacts Asp24 and Asn27 in H4 tail and Asp52 and Arg55 in H4 α2. **d)** Classification of Cryo-EM data of Oct4 bound nucleosome (Fig. 1a). The additional densities resembling interacting nucleosome ^37^ are differently positioned depending on Oct4 binding.

**Extended Data 5.**
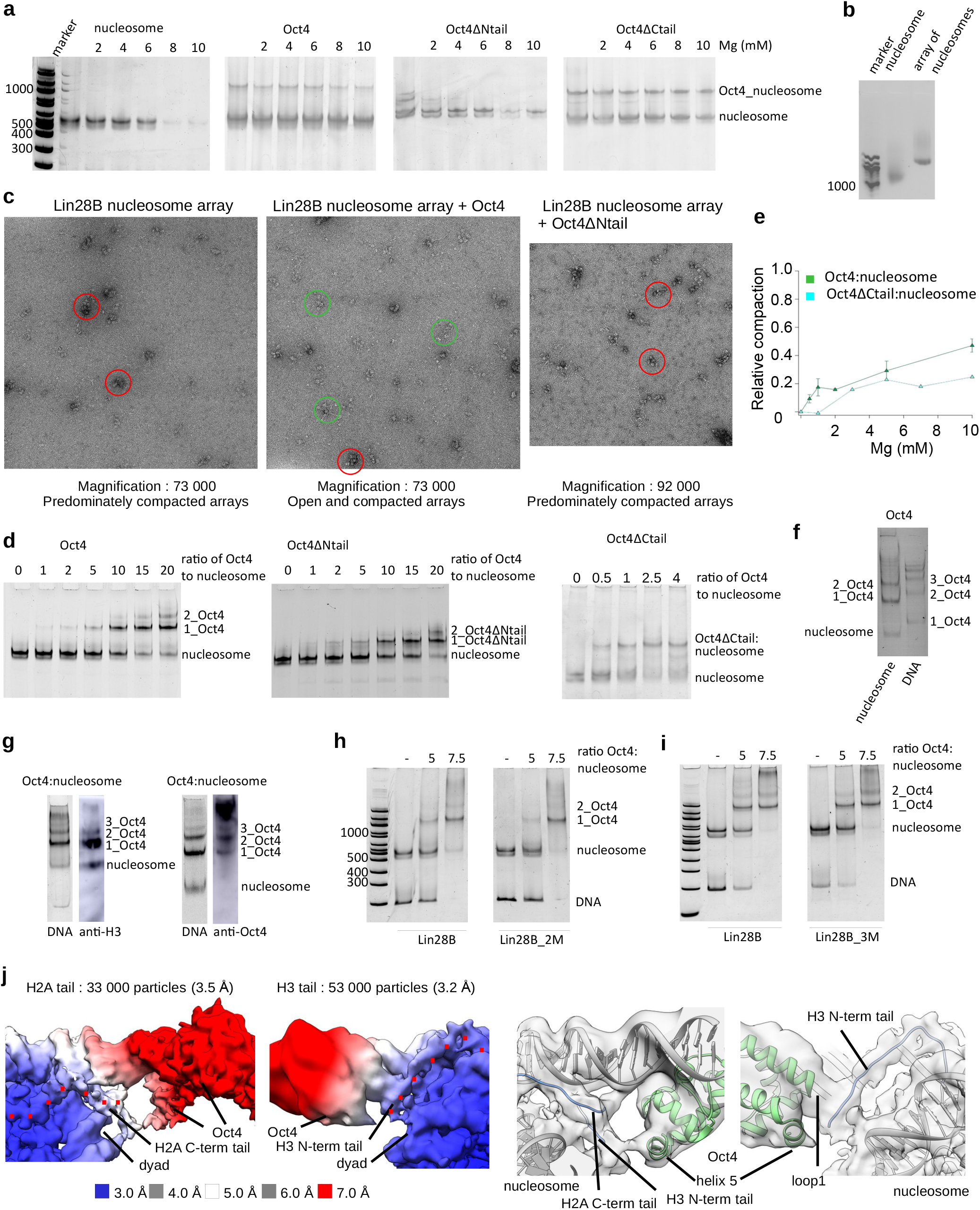
N-terminal disordered region of Oct4 is required for chromatin de-compaction. **a)** Native gels showing Mg^2+^ induced compaction of nucleosomes and nucleosomes bound to Oct4, Oct4ΔNtail and Oct4ΔCtail. Oct4 binding reduces nucleosome compaction. Deletion of Oct4 N-terminal disordered region eliminates Oct4 effect on nucleosome compaction. **b)** Native agarose gel showing assembly of nucleosome and nucleosome array. **c)** Negative stain micrographs showing Mg^2+^ induced compaction of the Lin28B nucleosome array, the Oct4 bound Lin28B nucleosome array and the Oct4ΔNtail bound Lin28B nucleosome array. Most nucleosomes are compacted (red circle) in sample containing Lin28B arrays and Oct4ΔNtail bound Lin28B arrays. Many more open arrays (green circle) are detectable when Oct4 is bound to Lin28B arrays. **d)** Native gels showing binding of Oct4ΔNtail and Oct4ΔCtail to the Lin28B nucleosome. Oct4ΔNtail and Oct4ΔCtail binds nucleosome comparably to wild type Oct4. **e)** Quantification of data from a). Deletion of Oct4 C-terminal disordered tail does not reduce Oct4 effect on nucleosome compaction. **f)** A native gel stained for DNA showing Oct4 binding to nucleosome and DNA. Binding to DNA and nucleosome generates distinct bands **g)** A native gel stained for DNA showing Oct4 binding to nucleosome and western blot with anti-H3 showing presence of histones in these complexes (left). A native gel stained for DNA showing Oct4 binding to nucleosome and western blot with anti-Oct4 showing presence of Oct4 in these complexes (right). **h)** Native gel showing Oct4 binding to the Lin28B nucleosome and Lin28B nucleosome with mutated binding site 2 (Lin28B_2M). Mutation of the binding site 2 did not affect binding of 1^st^ Oct4. **i)** Native gel showing Oct4 binding to the Lin28B nucleosome and Lin28B nucleosome with mutated binding site 3 (Lin28B_3M). Mutation of the binding site 3 did not affect binding of 1^st^ Oct4. One representative experiment of at least 3 independent experiments is shown for all biochemical data. **j)** Cryo-EM maps from a subset of data showing Oct4 interaction with H3 and H2A tails. The maps are colored by local resolution. Histone tails are marked with red dots. Model showing interaction of Oct4 with histone tails is shown below. Model of Oct4_nucleosome complex was rigid body fitted into cryoEM maps.

**Extended Data 6.**
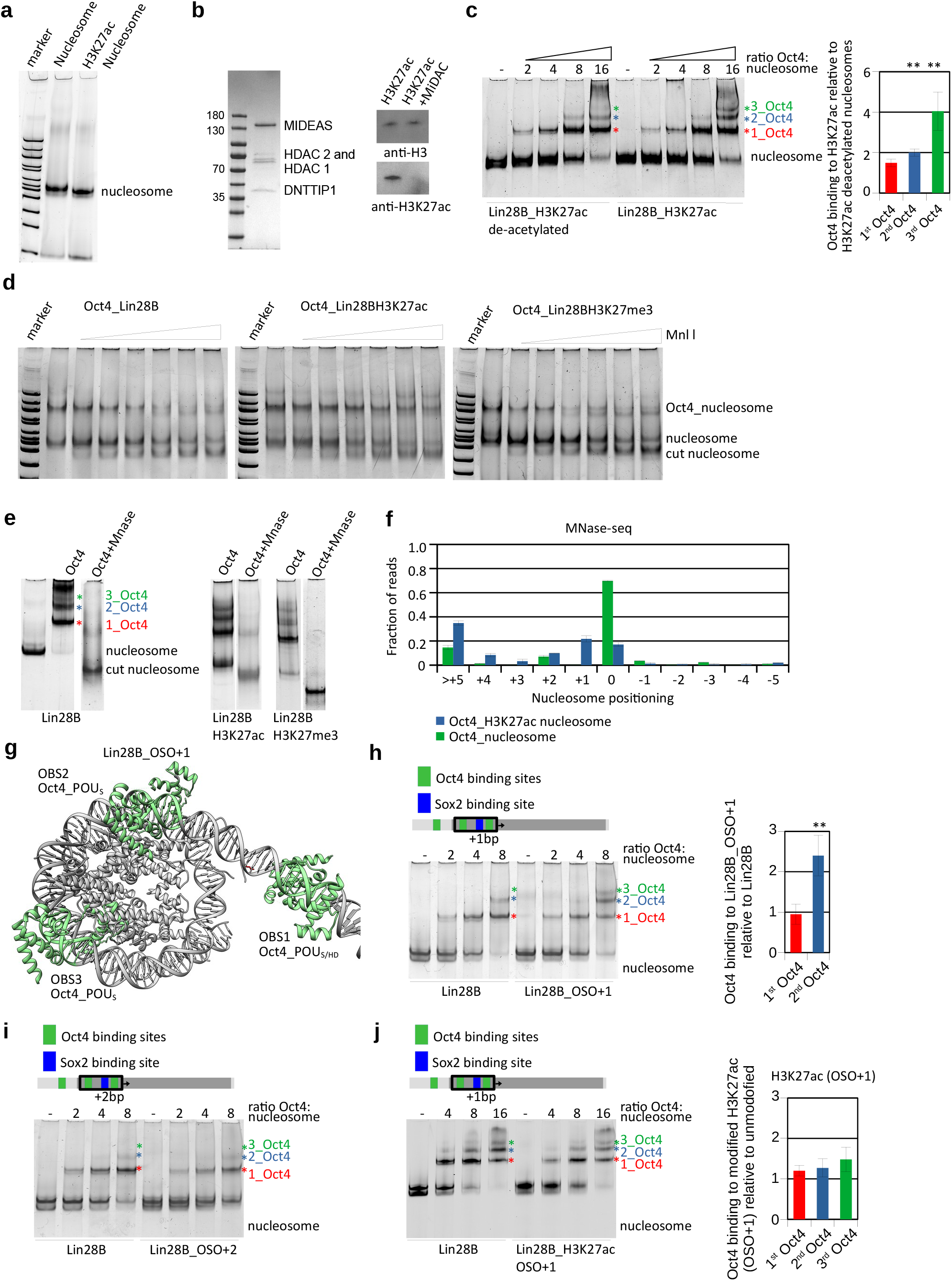
Histone modifications modulate Oct4 cooperativity. **a)** Native gel showing assembly of unmodified and H3K27ac Lin28B nucleosomes. **b)** Left, SDS-PAGE showing purification of MiDAC complex. Right, western blot with anti-H3 and anti-H3K27ac showing deacetylation of H3K27ac. **c)** Left, representative native gel electrophoresis showing Oct4 binding to the deacetylated Lin28B_H3K27ac or Lin28B_H3K27ac nucleosomes. The composition of the Oct4-bound bands was validated by Western blot (Extended Data Fig. 5g). Colored asterisks indicate the number of Oct4 molecules bound to the nucleosome: red, 1 Oct4; blue, 2 Oct4; and green, 3 Oct4. Right, quantification of the native gel electrophoresis data, using bands marked with asterisks. Data are mean and s.e.m. of 4 independent experiments; ** p<0.01, Student’s t-test. **d)** Mnl I digestion of unmodifed and H3K27ac nucleosomes bound to Oct4. Binding of Oct4 to nucleosomes increases Mnl I digestion of nucleosome indicating exposure of Mnl I site. Oct4 bound to H3K27ac nucleosomes shows decreased degradation at Mnl I site compared to unmodified nucleosomes bound to Oct4. **e)** Native gel showing Oct4 binding and MNase digestion of Oct4 bound unmodified and H3K27ac Lin28B nuclesomes. **f)** Quantification of sequencing of Mnase I digested Oct4-bound nucleosomes (unmodified and H3K27ac). The y-axis shows fraction of nucleosome size reads starting at defined position, the x-axis shows position of the first base pair relative to the most abundant position (0 as observed in the structure). Data are mean and s.e.m. of 2 independent experiments. **g)** Model of Oct4 bound to the Lin28B nucleosome with OBS2/3 and Sox2 binding sites moved for +1bp (OSO+1), showing Oct4-binding sites on DNA in green. Oct4 bound to OBS1 is in solid green; Oct4 structure was superimposed on OBS2 and on OBS3. In this conformation, Oct4_POU_S_ can bind to OBS2 and OBS3. Note, shift of 1bp exposes Oct4_POU_S_ binding site at OBS2 instead of Oct4_POU_HD_. **h)** Left, representative native gel electrophoresis showing Oct4 binding to the Lin28B or Lin28B nucleosomes with OBS2/3 and Sox2 binding sites moved for +1bp (OSO+1). Right, quantification of the native gel electrophoresis, data showns as s.e.m. of 3 independent experiments. Bands marked with ** were used for quantification. ** p<0.01. **i)** Representative native gel electrophoresis showing Oct4 binding to the Lin28B or Lin28B nucleosomes with OBS2/3 and Sox2 binding sites moved for +2bp (OSO+2). **j)** Left, representative native gel electrophoresis showing Oct4 binding to the Lin28B or H3K27ac Lin28B nucleosomes with OBS2/3 and Sox2 binding sites moved for +1bp (OSO+1). Right, quantification of the native gel electrophporesis, data showns as s.e.m. of 3 independent experiments. Bands marked with * were used for quantification.

**Extended Data 7.**
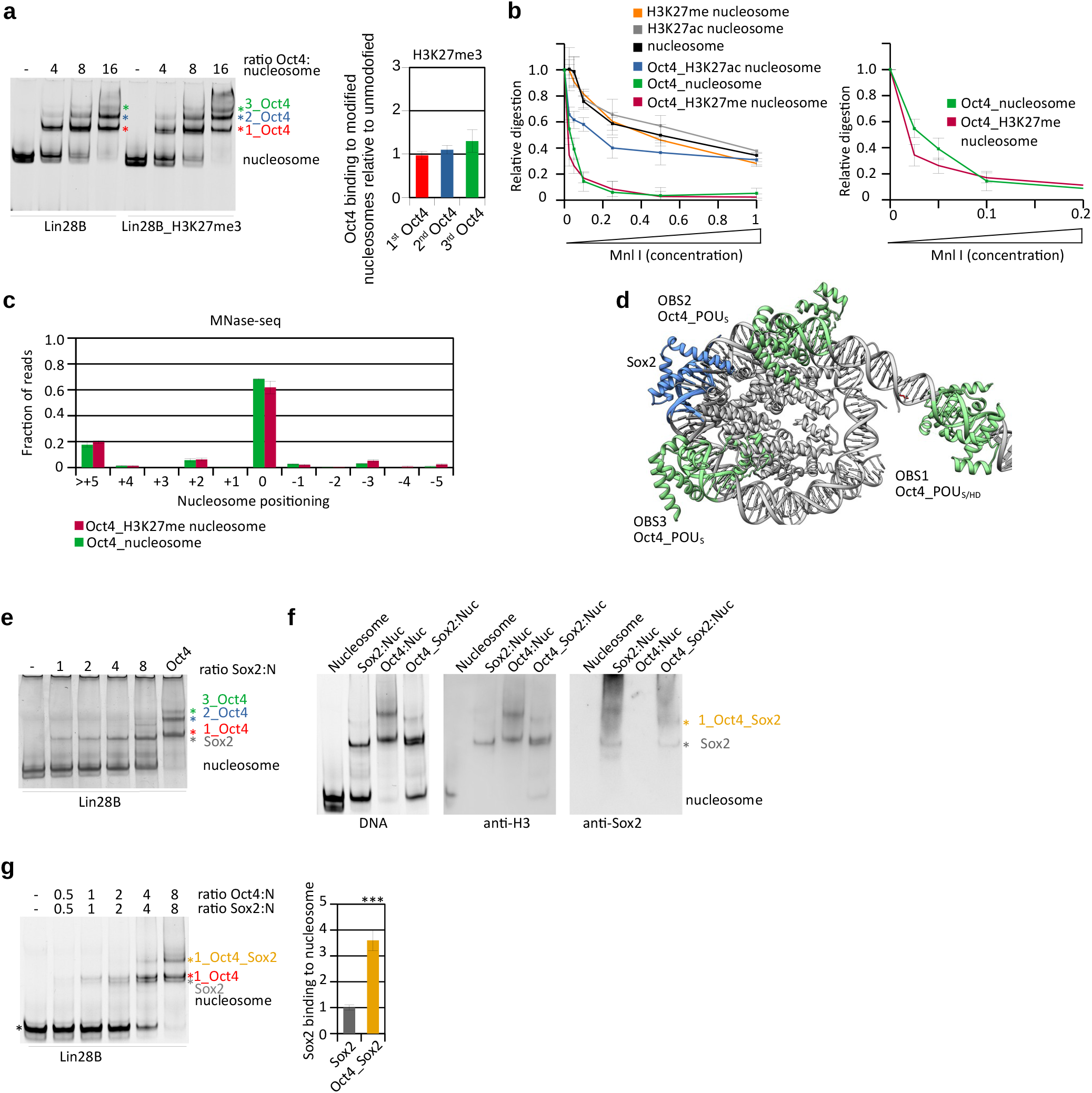
Histone modifications modulate Oct4 and Sox2 cooperativity. **a)** Left, representative native gel electrophoresis showing Oct4 binding to the Lin28B or H3K27me3 Lin28B nucleosomes. Right, quantification of the native gel electrophporesis, data showns as s.e.m. of 6 independent experiments. Bands marked with * were used for quantification. **b)** Left, quantification of Mnl I digestion of free and Oct4-bound nucleosomes (unmodified, H3K27ac and H3K27me3). The y-axis shows intensity of the intact Oct4_nucleosome complex or nucleosome band after enzyme digestion, normalized to the input (without enzyme), data are mean and s.e.m. of 4 independent experiments. Representative gels are shown in Extended Data Fig. 6d. Right, comparison of Mnl I digestion between unmodified and H3K27me3 Oct4 bound nucleosomes. **c)** Quantification of sequencing of Mnase I digested Oct4-bound nucleosomes (unmodified and H3K27me3). The y-axis shows fraction of nucleosome size reads starting at defined position, the x-axis shows position of the first base pair relative to the most abundant position (0 as observed in the structure). Data are mean and s.e.m. of 2 independent experiments. **d)** Model of Oct4 bound to the Lin28B nucleosome with Sox2 binding site shown in blue. Oct4 bound to OBS1 is in solid green; Oct4 structure was superimposed on OBS2 and on OBS3. Sox2 binding site and Sox2 model are shown in blue. **e)** A native gel stained for DNA showing Sox2 and Oct4 binding to nucleosome. **f)** A native gel stained for DNA showing Sox2 and Oct4 binding to nucleosome (left) and western blot with anti-H3 showing presence of histones in the complexes (center). Western blot with anti-Sox2 showing presence of Sox2 in these complexes is shown on the right. **g)** Left, representative native gel electrophoresis showing Oct4 and Sox2 binding to the Lin28B nucleosome. Oct4 and Sox2 were mixed and added to nucleosomes as indicated. Sox2 binding to nucleosome was validated by western blot analyses (Extended Data 7f). Colored asterisks indicate the molecules bound to the nucleosome: black, nucleosome alone; red, 1 Oct4; gray, 1 Sox2; orange, 1 Oct4 and 1 Sox2. Compare 1_Oct4_Sox2 band (orange asterisk) to 1_Oct4 band (red asterisk); and compare Sox2-bound nucleosome (gray asterisk) to input nucleosome (black asterisk). Right, quantification of the native gel electrophoresis data showing the described comparisons as ratios, using bands marked with asterisks. Data are mean and s.e.m. of 5 independent experiments; *** p<0.001, Student’s t-test.

**Extended Data 8.**
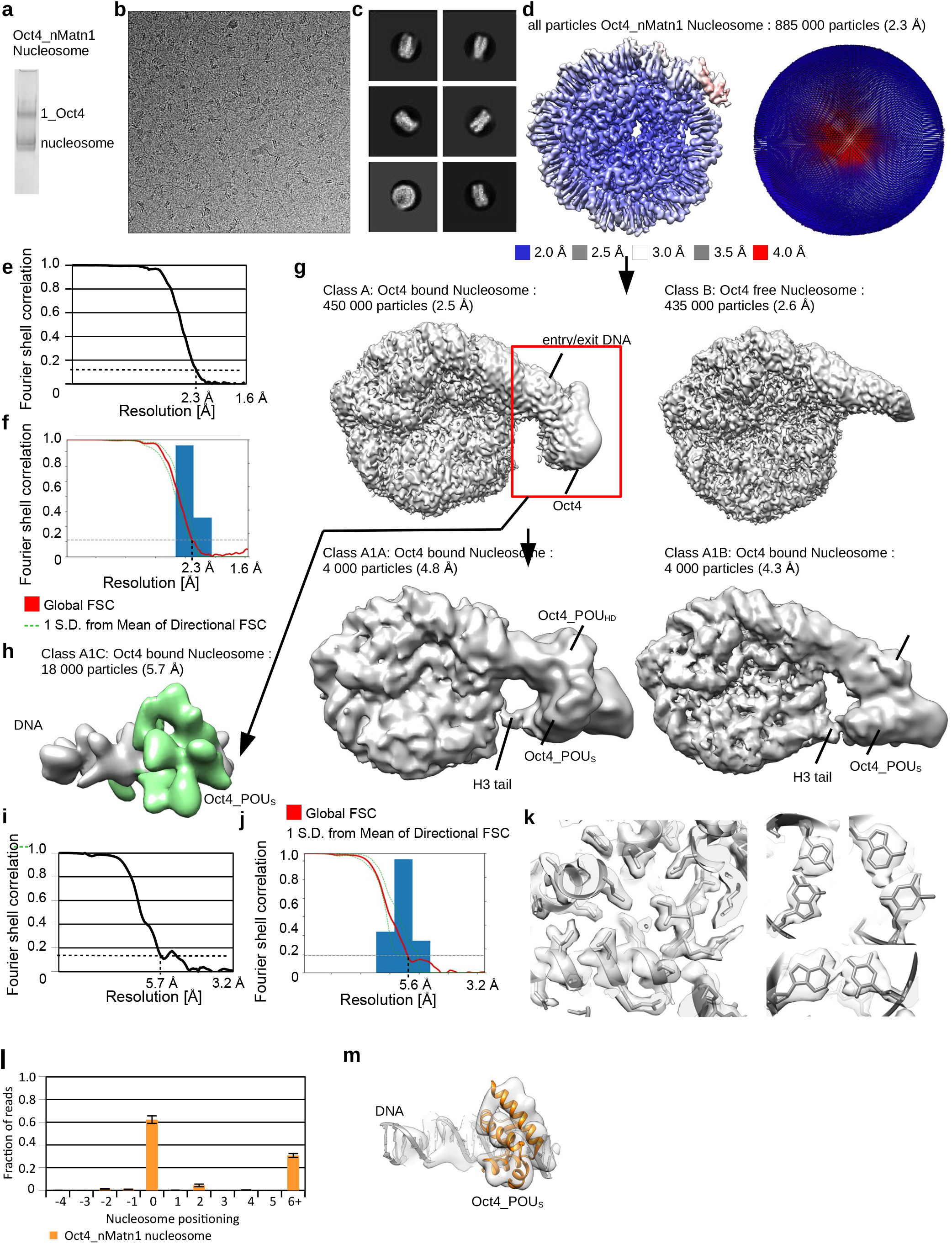
Cryo-EM of Oct4 bound to nMatn1 nucleosome. **a)** SDS-PAGE showing Oct4 binding to nMatn1 nucleosome. **b)** Representative cryo-EM micrograph collected with Titan Krios electron microscope at 300 keV. Nucleosome particles in multiple orientations are visible. **c)** Representative 2D class averages showing nucleosomes. Many details in nucleosomes are visible in 2D class averages. **d)** Cryo-EM map of nucleosome from the entire dataset, refined to 2.3 Å. The map is colored by local resolution. Angular distribution for nucleosome is shown on the right. **e)** Fourier shell correlation (FSC) curve showing the resolution of the map in d). **f)** Directional FSC plot showing uniform resolution in all directions. **g)** Classification of the nucleosome from d). Classification revealed two classes, nucleosome and nucleosome with bound Oct4. Oct4 bound class was further classified revealing many conformations of Oct4 bound to linker DNA. Two conformations with Oct4 bound to linker DNA are shown. Note Oct4 interaction with the H3 tail. **h)** Cryo-EM map of Oct4 region from the Oct4_nucleosome complex from g). Focused classification and refinements improved the resolution of this 25 kDa fragment to 5.6 Å. **i)** Fourier shell correlation (FSC) curve showing the resolution of the map in h). **j)** Directional FSC plot showing uniform resolution in all directions. **k)** Representative regions showing map quality and fit of the model are shown for the nucleosome with bound Oct4. Right: bases in the DNA are well resolved. **l)** Quantification of sequencing of Mnase I digested Oct4-bound nMatn1 nucleosomes. The y-axis shows fraction of nucleosome size reads starting at defined position, the x-axis shows position of the first base pair relative to the most abundant position (0 as observed in the structure). Data are mean and s.e.m. of 2 independent experiments. **m)** The model of the Oct4 bound to DNA (Extended Data 2g) was refined into the cryo-EM map. The representative region showing map quality and fit of the model is shown.

**Extended Data 9.**
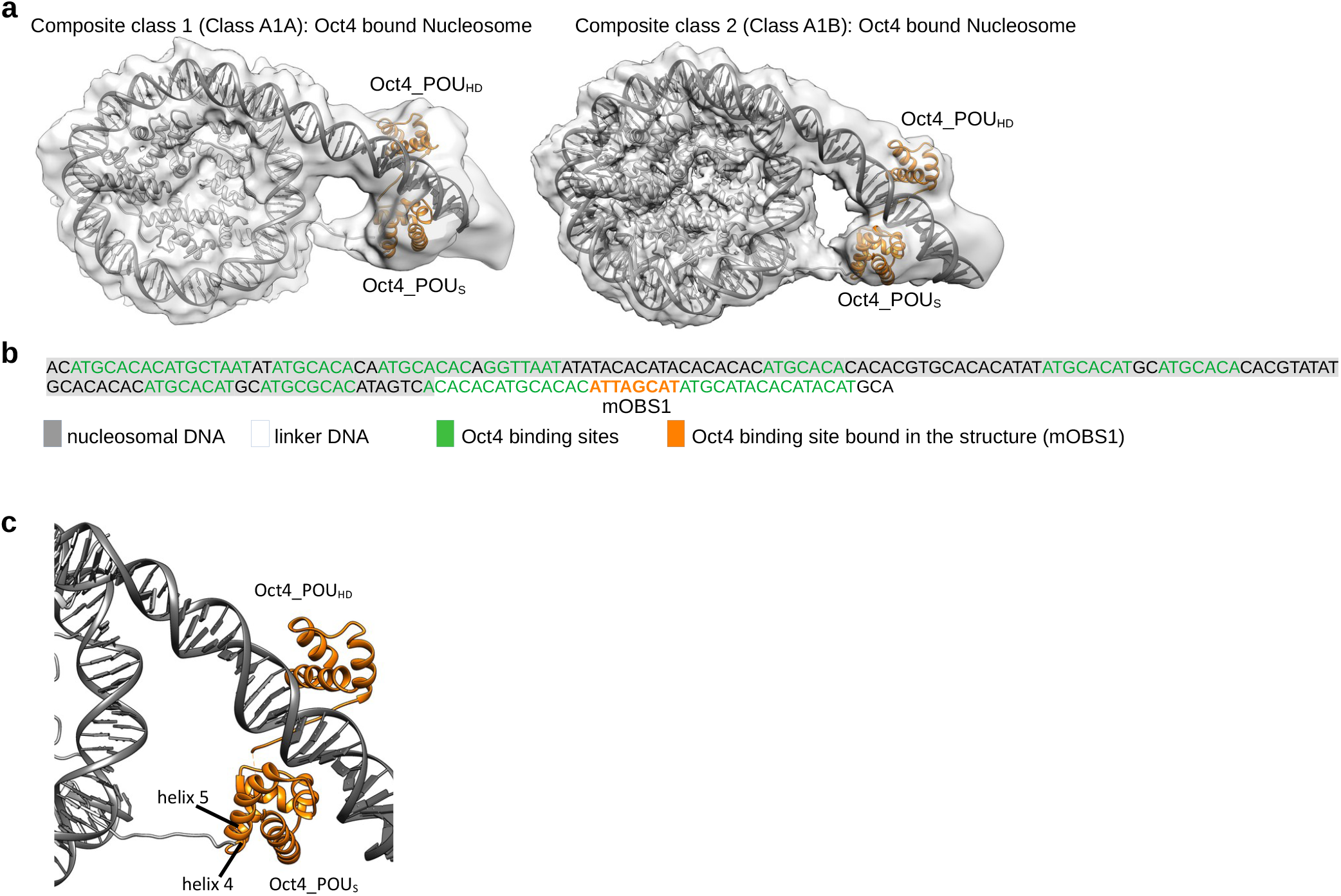
Histone modifications modulate Oct4 and Sox2 cooperativity on various human DNA. **a)** Cryo-EM models of Oct4 bound to the nMatn1 nucleosome containing 186bp of DNA at 2.2-5.6 Å resolution for two most dominant conformations. **b)** DNA sequence and schematic representation showing nMatn1 DNA positioning on the Oct4_nucleosome complex. Potential Oct4 binding sites are labeled in green. Oct4 binding site occupied in the structure is labeled in orange. **c)** Close-up views of the nucleosome entry/exit site showing interaction of the Oct4_POU_S_ domain with the H3 N-terminal tail. Ribbon representation shows Oct4_POU_S_ helix 4 and helix 5 interacting with histone H3 N-terminal tail.

**Extended Data 10.**
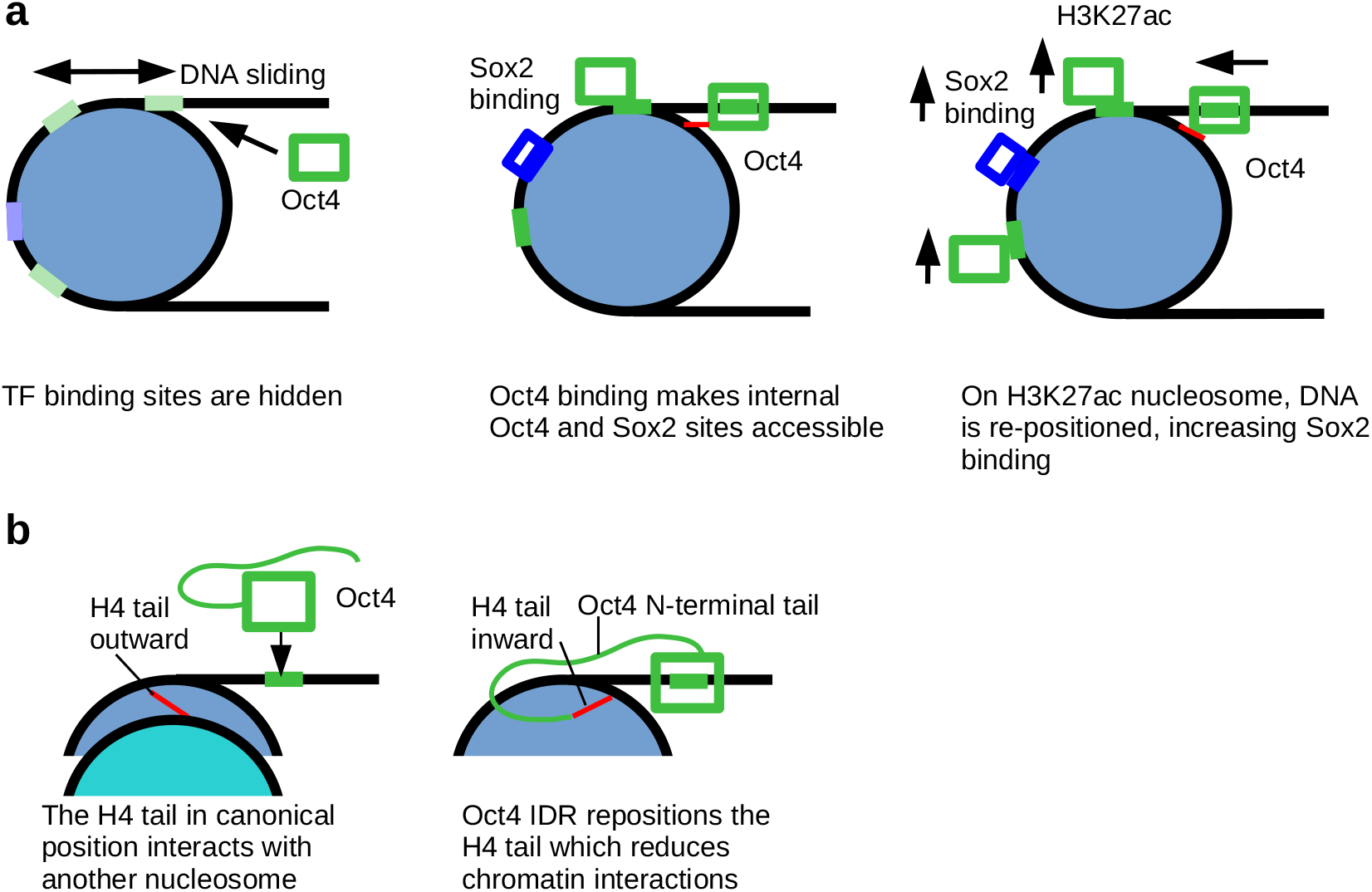
Model showing Oct4 binding to unmodified, H3K27ac and H3K27me3 nucleosomes. **a)** In Lin28B nucleosome Oct4 (light green) and Sox2 (light blue) binding sites are wrapped around the histone octamer. Lin28B nucleosomes are mobile, and nucleosome sliding transiently exposes the Oct4 binding site 1 (green), which leads to binding of Oct4 (green box). Oct4 binding (green box) traps DNA in a defined position, which positions and exposes internal Oct4 and Sox2 binding site (blue). Oct4 bound to the OBS1 interacts with histone H3 tail. H3K27ac abolishes this interaction leading to DNA movement towards the histone octamer, which exposes internal Oct4 and Sox2 sites even more, leading to increased binding. **b)** The canonical H4 tail conformation (red, facing outward) favors inter-nucleosome interactions by interacting with the acidic patch of neighboring nucleosomes. These interactions are essential for chromatin compaction. Oct4 DNA binding domain binds linker DNA whereas disordered activation domain binds H4 near the H4 tail. This repositions the H4 tail to an inward conformation that is incompatible with inter-nucleosome interactions in chromatin.

## References

1. Zaret, K.S. (2020). Pioneer Transcription Factors Initiating Gene Network Changes. Annu Rev Genet 54, 367–385. 10.1146/annurev-genet-030220-015007.

2. Iwafuchi-Doi, M., and Zaret, K.S. (2016). Cell fate control by pioneer transcription factors. Development 143, 1833–1837. 10.1242/dev.133900.

3. Lambert, S.A., Jolma, A., Campitelli, L.F., Das, P.K., Yin, Y., Albu, M., Chen, X., Taipale, J., Hughes, T.R., and Weirauch, M.T. (2018). The Human Transcription Factors. Cell 172, 650–665. 10.1016/j.cell.2018.01.029.

4. Cirillo, L.A., Lin, F.R., Cuesta, I., Friedman, D., Jarnik, M., and Zaret, K.S. (2002). Opening of compacted chromatin by early developmental transcription factors HNF3 (FoxA) and GATA-4. Mol Cell 9, 279–289. 10.1016/s1097-2765(02)00459-8.

5. Soufi, A., Donahue, G., and Zaret, K.S. (2012). Facilitators and impediments of the pluripotency reprogramming factors’ initial engagement with the genome. Cell 151, 994–1004. 10.1016/j.cell.2012.09.045.

6. Soufi, A., Garcia, M.F., Jaroszewicz, A., Osman, N., Pellegrini, M., and Zaret, K.S. (2015). Pioneer transcription factors target partial DNA motifs on nucleosomes to initiate reprogramming. Cell 161, 555–568. 10.1016/j.cell.2015.03.017.

7. Wang, J., Zhuang, J., Iyer, S., Lin, X., Whitfield, T.W., Greven, M.C., Pierce, B.G., Dong, X., Kundaje, A., Cheng, Y., et al. (2012). Sequence features and chromatin structure around the genomic regions bound by 119 human transcription factors. Genome Res 22, 1798–1812. 10.1101/gr.139105.112.

8. Zhu, F., Farnung, L., Kaasinen, E., Sahu, B., Yin, Y., Wei, B., Dodonova, S.O., Nitta, K.R., Morgunova, E., Taipale, M., et al. (2018). The interaction landscape between transcription factors and the nucleosome. Nature 562, 76–81. 10.1038/s41586-018-0549-5.

9. Takahashi, K., and Yamanaka, S. (2006). Induction of pluripotent stem cells from mouse embryonic and adult fibroblast cultures by defined factors. Cell 126, 663–676. 10.1016/j.cell.2006.07.024.

10. Kim, J.B., Sebastiano, V., Wu, G., Araúzo-Bravo, M.J., Sasse, P., Gentile, L., Ko, K., Ruau, D., Ehrich, M., van den Boom, D., et al. (2009). Oct4-induced pluripotency in adult neural stem cells. Cell 136, 411–419. 10.1016/j.cell.2009.01.023.

11. Wu, T., Wang, H., He, J., Kang, L., Jiang, Y., Liu, J., Zhang, Y., Kou, Z., Liu, L., Zhang, X., et al. (2011). Reprogramming of trophoblast stem cells into pluripotent stem cells by Oct4. Stem Cells 29, 755–763. 10.1002/stem.617.

12. Boyer, L.A., Lee, T.I., Cole, M.F., Johnstone, S.E., Levine, S.S., Zucker, J.P., Guenther, M.G., Kumar, R.M., Murray, H.L., Jenner, R.G., et al. (2005). Core transcriptional regulatory circuitry in human embryonic stem cells. Cell 122, 947–956. 10.1016/j.cell.2005.08.020.

13. King, H.W., and Klose, R.J. (2017). The pioneer factor OCT4 requires the chromatin remodeller BRG1 to support gene regulatory element function in mouse embryonic stem cells. Elife 6. 10.7554/eLife.22631.

14. Sönmezer, C., Kleinendorst, R., Imanci, D., Barzaghi, G., Villacorta, L., Schübeler, D., Benes, V., Molina, N., and Krebs, A.R. (2020). Molecular Co-occupancy Identifies Transcription Factor Binding Cooperativity In Vivo. Mol Cell. 10.1016/j.molcel.2020.11.015.

15. Tapia, N., MacCarthy, C., Esch, D., Gabriele Marthaler, A., Tiemann, U., Araúzo-Bravo, M.J., Jauch, R., Cojocaru, V., and Schöler, H.R. (2015). Dissecting the role of distinct OCT4-SOX2 heterodimer configurations in pluripotency. Sci Rep 5, 13533. 10.1038/srep13533.

16. Nichols, J., Zevnik, B., Anastassiadis, K., Niwa, H., Klewe-Nebenius, D., Chambers, I., Schöler, H., and Smith, A. (1998). Formation of pluripotent stem cells in the mammalian embryo depends on the POU transcription factor Oct4. Cell 95, 379–391. 10.1016/s0092-8674(00)81769-9.

17. Ambrosetti, D.C., Basilico, C., and Dailey, L. (1997). Synergistic activation of the fibroblast growth factor 4 enhancer by Sox2 and Oct-3 depends on protein-protein interactions facilitated by a specific spatial arrangement of factor binding sites. Mol Cell Biol 17, 6321–6329. 10.1128/MCB.17.11.6321.

18. Avilion, A.A., Nicolis, S.K., Pevny, L.H., Perez, L., Vivian, N., and Lovell-Badge, R. (2003). Multipotent cell lineages in early mouse development depend on SOX2 function. Genes Dev 17, 126–140. 10.1101/gad.224503.

19. Esch, D., Vahokoski, J., Groves, M.R., Pogenberg, V., Cojocaru, V., Vom Bruch, H., Han, D., Drexler, H.C.A., Araúzo-Bravo, M.J., Ng, C.K.L., et al. (2013). A unique Oct4 interface is crucial for reprogramming to pluripotency. Nat Cell Biol 15, 295–301. 10.1038/ncb2680.

20. Reményi, A., Lins, K., Nissen, L.J., Reinbold, R., Schöler, H.R., and Wilmanns, M. (2003). Crystal structure of a POU/HMG/DNA ternary complex suggests differential assembly of Oct4 and Sox2 on two enhancers. Genes Dev 17, 2048–2059. 10.1101/gad.269303.

21. Michael, A.K., Grand, R.S., Isbel, L., Cavadini, S., Kozicka, Z., Kempf, G., Bunker, R.D., Schenk, A.D., Graff-Meyer, A., Pathare, G.R., et al. (2020). Mechanisms of OCT4-SOX2 motif readout on nucleosomes. Science 368, 1460–1465. 10.1126/science.abb0074.

22. Lowary, P.T., and Widom, J. (1998). New DNA sequence rules for high affinity binding to histone octamer and sequence-directed nucleosome positioning. J. Mol. Biol. 276, 19–42. 10.1006/jmbi.1997.1494.

23. Roberts, G.A., Ozkan, B., Gachulincová, I., O’Dwyer, M.R., Hall-Ponsele, E., Saxena, M., Robinson, P.J., and Soufi, A. (2021). Dissecting OCT4 defines the role of nucleosome binding in pluripotency. Nat Cell Biol 23, 834–845. 10.1038/s41556-021-00727-5.

24. Echigoya, K., Koyama, M., Negishi, L., Takizawa, Y., Mizukami, Y., Shimabayashi, H., Kuroda, A., and Kurumizaka, H. (2020). Nucleosome binding by the pioneer transcription factor OCT4. Sci Rep 10, 11832. 10.1038/s41598-020-68850-1.

25. Iurlaro, M., Stadler, M.B., Masoni, F., Jagani, Z., Galli, G.G., and Schübeler, D. (2021). Mammalian SWI/SNF continuously restores local accessibility to chromatin. Nat Genet 53, 279–287. 10.1038/s41588-020-00768-w.

26. Chen, J., Chen, X., Li, M., Liu, X., Gao, Y., Kou, X., Zhao, Y., Zheng, W., Zhang, X., Huo, Y., et al. (2016). Hierarchical Oct4 Binding in Concert with Primed Epigenetic Rearrangements during Somatic Cell Reprogramming. Cell Rep 14, 1540–1554. 10.1016/j.celrep.2016.01.013.

27. Donaghey, J., Thakurela, S., Charlton, J., Chen, J.S., Smith, Z.D., Gu, H., Pop, R., Clement, K., Stamenova, E.K., Karnik, R., et al. (2018). Genetic determinants and epigenetic effects of pioneer-factor occupancy. Nat Genet 50, 250–258. 10.1038/s41588-017-0034-3.

28. Lee, T.I., Jenner, R.G., Boyer, L.A., Guenther, M.G., Levine, S.S., Kumar, R.M., Chevalier, B., Johnstone, S.E., Cole, M.F., Isono, K., et al. (2006). Control of developmental regulators by Polycomb in human embryonic stem cells. Cell 125, 301–313. 10.1016/j.cell.2006.02.043.

29. Benveniste, D., Sonntag, H.-J., Sanguinetti, G., and Sproul, D. (2014). Transcription factor binding predicts histone modifications in human cell lines. Proc Natl Acad Sci U S A 111, 13367–13372. 10.1073/pnas.1412081111.

30. Kim, J., Chu, J., Shen, X., Wang, J., and Orkin, S.H. (2008). An extended transcriptional network for pluripotency of embryonic stem cells. Cell 132, 1049–1061. 10.1016/j.cell.2008.02.039.

31. Hanna, J., Saha, K., Pando, B., van Zon, J., Lengner, C.J., Creyghton, M.P., van Oudenaarden, A., and Jaenisch, R. (2009). Direct cell reprogramming is a stochastic process amenable to acceleration. Nature 462, 595–601. 10.1038/nature08592.

32. Tsialikas, J., and Romer-Seibert, J. (2015). LIN28: roles and regulation in development and beyond. Development 142, 2397–2404. 10.1242/dev.117580.

33. Yu, J., Vodyanik, M.A., Smuga-Otto, K., Antosiewicz-Bourget, J., Frane, J.L., Tian, S., Nie, J., Jonsdottir, G.A., Ruotti, V., Stewart, R., et al. (2007). Induced pluripotent stem cell lines derived from human somatic cells. Science 318, 1917–1920. 10.1126/science.1151526.

34. Kelly, T.K., Liu, Y., Lay, F.D., Liang, G., Berman, B.P., and Jones, P.A. (2012). Genome-wide mapping of nucleosome positioning and DNA methylation within individual DNA molecules. Genome Res 22, 2497–2506. 10.1101/gr.143008.112.

35. Bilokapic, S., Strauss, M., and Halic, M. (2018). Histone octamer rearranges to adapt to DNA unwrapping. Nat. Struct. Mol. Biol. 25, 101–108. 10.1038/s41594-017-0005-5.

36. Luger, K., Mäder, A.W., Richmond, R.K., Sargent, D.F., and Richmond, T.J. (1997). Crystal structure of the nucleosome core particle at 2.8 A resolution. Nature 389, 251–260. 10.1038/38444.

37. Bilokapic, S., Strauss, M., and Halic, M. (2018). Cryo-EM of nucleosome core particle interactions in trans. Sci Rep 8, 7046. 10.1038/s41598-018-25429-1.

38. de Frutos, M., Raspaud, E., Leforestier, A., and Livolant, F. (2001). Aggregation of nucleosomes by divalent cations. Biophys J 81, 1127–1132. 10.1016/S0006-3495(01)75769-4.

39. McBryant, S.J., Klonoski, J., Sorensen, T.C., Norskog, S.S., Williams, S., Resch, M.G., Toombs, J.A., Hobdey, S.E., and Hansen, J.C. (2009). Determinants of histone H4 N-terminal domain function during nucleosomal array oligomerization: roles of amino acid sequence, domain length, and charge density. J Biol Chem 284, 16716–16722. 10.1074/jbc.M109.011288.

40. Lowary, P.T., and Widom, J. (1997). Nucleosome packaging and nucleosome positioning of genomic DNA. Proc. Natl. Acad. Sci. U.S.A. 94, 1183–1188.

41. Meers, M.P., Janssens, D.H., and Henikoff, S. (2019). Pioneer Factor-Nucleosome Binding Events during Differentiation Are Motif Encoded. Mol Cell 75, 562–575.e5. 10.1016/j.molcel.2019.05.025.

42. Judd, J., Duarte, F.M., and Lis, J.T. (2021). Pioneer-like factor GAF cooperates with PBAP (SWI/SNF) and NURF (ISWI) to regulate transcription. Genes Dev 35, 147–156. 10.1101/gad.341768.120.

43. Pardo, M., Lang, B., Yu, L., Prosser, H., Bradley, A., Babu, M.M., and Choudhary, J. (2010). An expanded Oct4 interaction network: implications for stem cell biology, development, and disease. Cell Stem Cell 6, 382–395. 10.1016/j.stem.2010.03.004.

44. Swinstead, E.E., Paakinaho, V., Presman, D.M., and Hager, G.L. (2016). Pioneer factors and ATP-dependent chromatin remodeling factors interact dynamically: A new perspective: Multiple transcription factors can effect chromatin pioneer functions through dynamic interactions with ATP-dependent chromatin remodeling factors. Bioessays 38, 1150–1157. 10.1002/bies.201600137.

45. Vierbuchen, T., Ling, E., Cowley, C.J., Couch, C.H., Wang, X., Harmin, D.A., Roberts, C.W.M., and Greenberg, M.E. (2017). AP-1 Transcription Factors and the BAF Complex Mediate Signal-Dependent Enhancer Selection. Mol Cell 68, 1067–1082.e12. 10.1016/j.molcel.2017.11.026.

46. Wang, L., Du, Y., Ward, J.M., Shimbo, T., Lackford, B., Zheng, X., Miao, Y., Zhou, B., Han, L., Fargo, D.C., et al. (2014). INO80 facilitates pluripotency gene activation in embryonic stem cell self-renewal, reprogramming, and blastocyst development. Cell Stem Cell 14, 575–591. 10.1016/j.stem.2014.02.013.

47. Dodonova, S.O., Zhu, F., Dienemann, C., Taipale, J., and Cramer, P. (2020). Nucleosome-bound SOX2 and SOX11 structures elucidate pioneer factor function. Nature 580, 669–672. 10.1038/s41586-020-2195-y.

48. Iwafuchi, M., Cuesta, I., Donahue, G., Takenaka, N., Osipovich, A.B., Magnuson, M.A., Roder, H., Seeholzer, S.H., Santisteban, P., and Zaret, K.S. (2020). Gene network transitions in embryos depend upon interactions between a pioneer transcription factor and core histones. Nat Genet 52, 418–427. 10.1038/s41588-020-0591-8.

49. Luger, K., Rechsteiner, T.J., and Richmond, T.J. (1999). Expression and purification of recombinant histones and nucleosome reconstitution. Methods Mol. Biol. 119, 1–16. 10.1385/1-59259-681-9:1.

50. Ivic N, Groschup B, Bilokapic S, and Halic M (2016). Simplified Method for Rapid Purification of Soluble Histones. Croatica chemica acta 89, 153–162.

51. Bilokapic, S., and Halic, M. (2019). Nucleosome and ubiquitin position Set2 to methylate H3K36. Nat Commun 10, 3795. 10.1038/s41467-019-11726-4.

52. Bilokapic, S., Strauss, M., and Halic, M. (2018). Structural rearrangements of the histone octamer translocate DNA. Nat Commun 9, 1330. 10.1038/s41467-018-03677-z.

53. Simon, M.D., Chu, F., Racki, L.R., de la Cruz, C.C., Burlingame, A.L., Panning, B., Narlikar, G.J., and Shokat, K.M. (2007). The site-specific installation of methyl-lysine analogs into recombinant histones. Cell 128, 1003–1012. 10.1016/j.cell.2006.12.041.

54. Carey, M.F., Peterson, C.L., and Smale, S.T. (2009). Dignam and Roeder nuclear extract preparation. Cold Spring Harb Protoc 2009, pdb.prot5330. 10.1101/pdb.prot5330.

55. Schneider, C.A., Rasband, W.S., and Eliceiri, K.W. (2012). NIH Image to ImageJ: 25 years of image analysis. Nat Methods 9, 671–675. 10.1038/nmeth.2089.

56. Grant, T., and Grigorieff, N. (2015). Measuring the optimal exposure for single particle cryoEM using a 2.6 Å reconstruction of rotavirus VP6. Elife 4, e06980. 10.7554/eLife.06980.

57. Zheng, S.Q., Palovcak, E., Armache, J.-P., Verba, K.A., Cheng, Y., and Agard, D.A. (2017). MotionCor2: anisotropic correction of beam-induced motion for improved cryo-electron microscopy. Nat. Methods 14, 331–332. 10.1038/nmeth.4193.

58. Rohou, A., and Grigorieff, N. (2015). CTFFIND4: Fast and accurate defocus estimation from electron micrographs. J. Struct. Biol. 192, 216–221. 10.1016/j.jsb.2015.08.008.

59. Wagner, T., Merino, F., Stabrin, M., Moriya, T., Antoni, C., Apelbaum, A., Hagel, P., Sitsel, O., Raisch, T., Prumbaum, D., et al. (2019). SPHIRE-crYOLO is a fast and accurate fully automated particle picker for cryo-EM. Commun Biol 2, 218. 10.1038/s42003-019-0437-z.

60. Zivanov, J., Nakane, T., Forsberg, B.O., Kimanius, D., Hagen, W.J., Lindahl, E., and Scheres, S.H. (2018). New tools for automated high-resolution cryo-EM structure determination in RELION-3. Elife 7. 10.7554/eLife.42166.

61. Terwilliger, T.C., Ludtke, S.J., Read, R.J., Adams, P.D., and Afonine, P.V. (2020). Improvement of cryo-EM maps by density modification. Nat Methods 17, 923–927. 10.1038/s41592-020-0914-9.

62. Emsley, P., Lohkamp, B., Scott, W.G., and Cowtan, K. (2010). Features and development of Coot. Acta Crystallogr. D Biol. Crystallogr. 66, 486–501. 10.1107/S0907444910007493.

63. Bilokapic, S., Suskiewicz, M.J., Ahel, I., and Halic, M. (2020). Bridging of DNA breaks activates PARP2-HPF1 to modify chromatin. Nature 585, 609–613. 10.1038/s41586-020-2725-7.

64. Adams, P.D., Afonine, P.V., Bunkóczi, G., Chen, V.B., Davis, I.W., Echols, N., Headd, J.J., Hung, L.-W., Kapral, G.J., Grosse-Kunstleve, R.W., et al. (2010). PHENIX: a comprehensive Pythonbased system for macromolecular structure solution. Acta Crystallogr. D Biol. Crystallogr. 66, 213–221. 10.1107/S0907444909052925.

65. Pettersen, E.F., Goddard, T.D., Huang, C.C., Couch, G.S., Greenblatt, D.M., Meng, E.C., and Ferrin, T.E. (2004). UCSF Chimera--a visualization system for exploratory research and analysis. J Comput Chem 25, 1605–1612. 10.1002/jcc.20084.

